# Active Transcription and Epigenetic Reactions Synergistically Regulate Meso-Scale Genomic Organization

**DOI:** 10.1101/2023.04.21.537822

**Authors:** Aayush Kant, Zixian Guo, Vinayak, Maria Victoria Neguembor, Wing Shun Li, Vasundhara Agrawal, Emily Pujadas, Luay Almassalha, Vadim Backman, Melike Lakadamyali, Maria Pia Cosma, Vivek B. Shenoy

## Abstract

In interphase nuclei, chromatin is organized into interspersed dense domains with characteristic sizes, both in the nuclear interior and periphery. However, the quantitative impact of transcription and histone modifications on the size and distribution of these domains remains unclear. Here, we introduce a mesoscale theoretical model that investigates the relationship between heterochromatic domain sizes and loop extrusion rates from these domains. The model considers chromatin-chromatin and chromatin-lamina interactions, methylation and acetylation kinetics, and diffusion of epigenetic marks and nucleoplasm. Our model generates testable predictions that help reveal the biophysics underlying chromatin organization in the presence of transcription-driven loop extrusion. This process is kinetically captured through the conversion of heterochromatin to euchromatin in response to RNAPII activity. We discovered that a balance between diffusive and reactive fluxes governs the steady-state sizes of heterochromatin domains. Using theory and simulations, we predicted that a loss of transcription results in increased chromatin compaction and larger heterochromatin domain sizes. To validate our predictions, we employed complementary super-resolution and nano-imaging techniques on five different cell lines with impaired transcription. We quantitatively assessed how domain sizes scale with loop extrusion rates at the hetero-euchromatin interfaces. Our analysis of previously obtained super-resolution images of nuclei revealed that excessive loop extrusion leads to smaller heterochromatin domains. The model successfully recapitulated these observations, explaining how transcription loss can counteract the effects of cohesin overloading. As the general biophysical mechanisms regulating heterochromatin domain sizes are independent of cell type, our findings have significant implications for understanding the role of transcription in global genome organization.

## 1 Introduction

The three-dimensional organization of chromatin within the nucleus is key to understanding the biophysical origin of critical cellular activities ranging from cell fate decisions to migration, proliferation, and metabolism. The existence of a multiscale chromatin organization is well-established [1, 2]. At the microscale, chromatin is organized into transcriptionally distinct compartments – a transcriptionally active, loosely packed euchromatin phase and a tightly packed, predominantly silent heterochromatin phase. Finer resolution of the chromatin conformation reveals the existence of a more detailed spatial organization ranging from self-interacting topologically associated domains (TADs) to chromatin loops bound by the CCCTC-binding factor (CTCF) and the cohesin complex [1, 2]. The chromatin fibers trapped in cohesin rings are extruded until either CTCF bound sites are encountered or cohesin is unloaded [2-4]. In addition to direct extrusion of DNA loops via cohesin motor activity [5-7], RNA polymerase II (RNAP II), a protein complex essential for DNA transcription, has been identified to play a significant role in enabling the movement of the chromatin fiber resulting in chromatin loop extrusion through cohesin [4, 8-14]. Specifically, the supercoiling of DNA due to transcriptional activity has been proposed to play a role in *in-vivo* chromatin loop extrusion [14, 15]. A recent experimental study combined super-resolution imaging of chromatin and single-molecule tracking of cohesin with various biological perturbations, such as pharmacological and genetic inhibition of transcription, supercoiling, and loop extrusion. This approach provided compelling evidence that transcription-mediated supercoiling regulates loop extrusion, as well as the spatial organization of chromatin within the nucleus [14]. These observations present a novel avenue of crosstalk between chromatin’s multiscale structural organization and its transcriptional status. This indicates that a bi-directional coupling exists, such that not only do the distinct phases of chromatin organization regulate transcription, but transcriptional activity can also affect genome organization via chromatin tethering, extrusion, and decompaction [14, 16]. While the local microscopic effects of transcription on spatial DNA organization have been previously investigated, a fundamental quantitative understanding of the physical mechanisms involved in the global genomic organization, due to transcriptional and epigenetic regulation, is not yet fully understood.

Here, we propose a novel, mesoscale coarse-grained, polymer physics-based mathematical model to capture the formation of chromatin domains while incorporating the spatio-temporal role of transcription-driven chromatin extrusion kinetics. Chromatin-chromatin interactions establish an energy landscape which drives a separation of hetero-and eu-chromatin phases through diffusion of the nucleoplasm and epigenetic marks. This process leads to the formation of functionally distinct heterochromatin domains of characteristic sizes. Chromatin-lamina interactions along the nuclear periphery give rise to lamina associated heterochromatin domains. The chromatin loop extrusion through active transcription is captured via the conversion of inactive heterochromatin into transcriptionally active euchromatin loops along the chromatin phase boundaries. Essential and unique to our model is the interplay of the epigenetic and transcriptional kinetics in governing meso-scale chromatin organization – including the size of heterochromatin domains and their spacing in the interior and periphery of the nucleus.

Using this model, we make quantitative predictions that offer a mechanistic explanation for the emergence of size scaling of compacted heterochromatin domains with the rate of loop extrusion at the domain interfaces. Importantly, by including the interactions of chromatin with the nuclear lamina, we show the quantitative dependence of the sizes of lamin-associated domains (LADs) as well as those of interior chromatin domains on the level of transcriptional activity. The predictions on the size scaling of heterochromatin domains made by the model are agnostic to specific interactions, and thus are not limited to a particular cell type. Indeed, the model predictions are qualitatively validated experimentally on five different cell lines and using two different nanoscopic imaging approaches. We used partial wave spectroscopy (PWS), which enables high-throughput, label-free, live cell imaging, in conjunction with scanning transmission electron microscopy tomography with ChromEM staining (ChromSTEM), which allows 3-dimensional high-resolution quantification of chromatin mass distribution, to quantify statistical domain properties upon inhibition of transcription. We, further, quantitatively validated our predictions by analyzing the length scales of compacted chromatin domains previously reported using stochastic optical reconstruction microscopy (STORM) imaging [14]. In conjunction with super-resolution microscopy and nano-imaging techniques, our model establishes a foundation for a predictive framework with broad implications for understanding the role of transcriptional and epigenetic crosstalk in defining mesoscale genome organization.

## 2 Materials and Methods

### 2.1 Mathematical description of genomic organization in the nucleus

At the meso-scale, chromatin is organized into distinct transcriptionally dissimilar phases of euchromatin and heterochromatin (Figure 1a). We treat the meso-scale genomic organization as a dynamic, far-from-equilibrium process, governed by the energetics of phase-separation in conjunction with the kinetics of epigenetic reactions and the formation of chromatin loops aided by DNA extrusion through cohesin due to RNAPII-mediated transcription. As shown in Figure 1a, the model considers energetics arising from chromatin-chromatin interactions leading to phase-separation and chromatin-lamina interactions leading to LAD formation. Time-dependent kinetics is regulated by (a) the free energy lowering diffusion of nucleoplasm and epigenetic marks, and (b) the active interconversion between the eu-and heterochromatic phases of chromatin. The active interconversion can take place in two ways: histone methylation or acetylation reactions can change the epigenetic distribution or active chromatin loop extrusion can drive the formation of euchromatin from heterochromatin.

**Figure 1.**
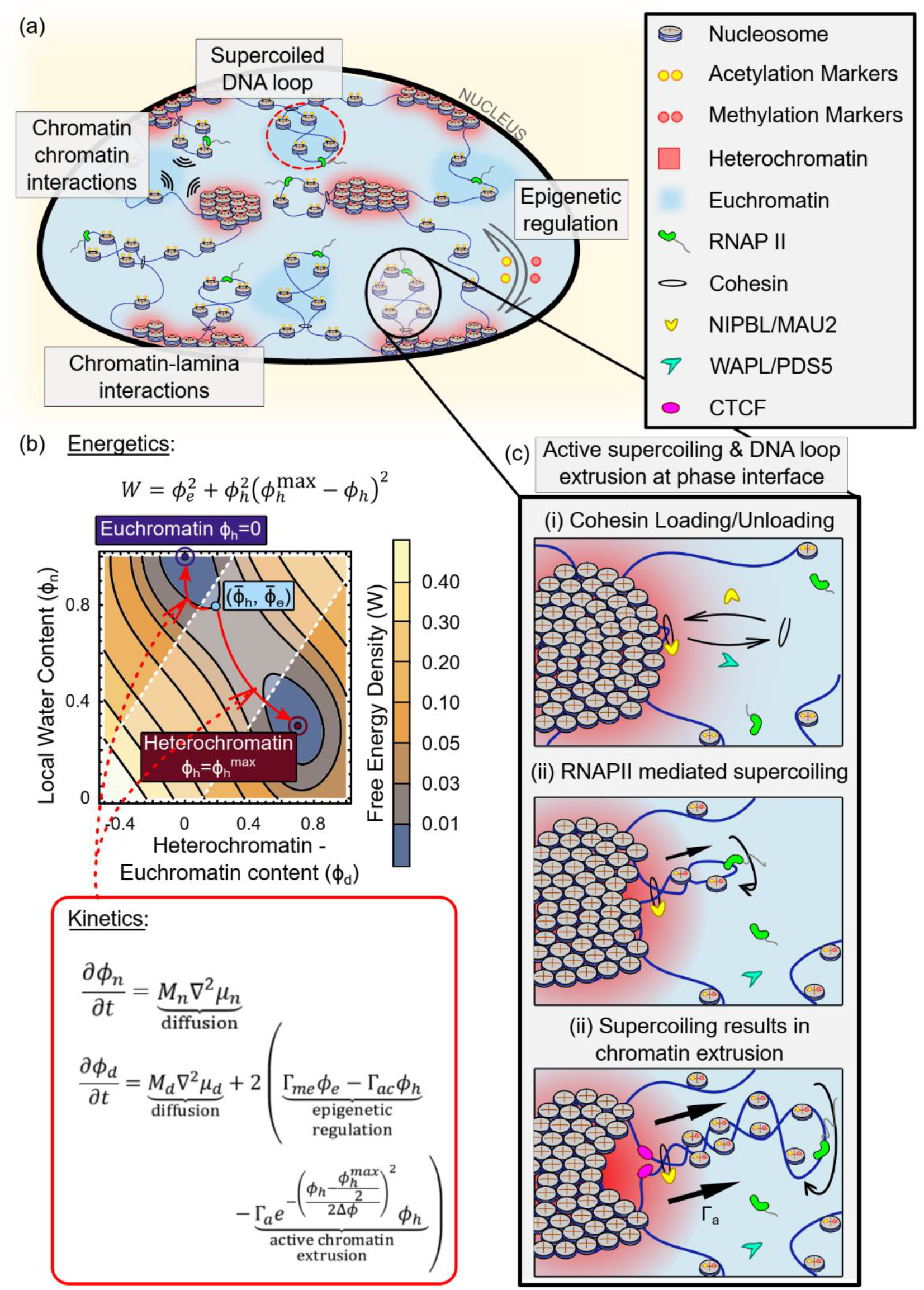
: (a) Schematic of a nucleus showing the multiple mechanisms involved in chromatin organization such as chromatin-chromatin interactions, the chromatin-lamina interactions and epigenetic regulation. Additionally, extrusion of chromatin loops due to DNA supercoiling – which is increased by transcriptional activity – also plays a role in meso-scale genomic organization. While this may occur within the chromatin phases (red circle), we further explore the role of chromatin loop extrusion at the heterochromatin-euchromatin interface (black circle). (b) The model captures the chromatin-chromatin interaction <u>energetics</u> via a double well free energy description as shown in the contour plot. The two wells correspond to the heterochromatin (red dot) and euchromatin phases (blue dot). Any initial configurations (light blue circle) spontaneously decompose into these wells at steady state. The dynamics of this transition is governed by diffusion and reaction <u>kinetics</u> comprising of epigenetic regulation and kinetics of active chromatin extrusion (red box inset). (c) Loading of cohesin assisted by NIPBL/MAU2 initiates the formation of chromatin loops. Cohesin can also be dynamically unloaded via unloading factors viz. WAPL/PDS5. Active processes such as RNAPII mediated transcription further drive the extrusion of trapped DNA, supercoiling it into chromatin loops.

As shown in Figure 1a, such RNAPII mediated DNA loop extrusion, can occur broadly in two regions: within the euchromatin domains (red dashed circle in Figure 1a) or at the interface of heterochromatin and euchromatin phases (black circle in Figure 1a). Since the extrusion of DNA loops in the euchromatin phase maintains its transcriptionally active status and does not lead to any significant mesoscale changes in the epigenetic distribution, we focus on the DNA loop extrusion at the domain interface. The DNA loop extrusion at the interface is instrumental in the reduction in size of heterochromatic domains at the periphery to form euchromatin.

### 2.2 Free energy considerations for the hetero- and eu-chromatic phases

We consider three nuclear constituents, namely the nucleoplasm and the two phases of chromatin, euchromatin and heterochromatin. We assume that at any point ***x*** in the nucleus, at a time *t*, these three constituents are space filling, and their volume fractions add up to unity, i.e., *ϕ*_*e*_ + *ϕ*_*h*_ + *ϕ*_*n*_ = 1. Hence, if the volume fractions of two of the constituents is known, the volume fraction of the third is determined by this constraint. The composition of the constituents can thus be defined in terms of two independent variables (refer to the methods for details) – (i) *ϕ*_*n*_(***x***, *t*) volume fraction of the <u>n</u>ucleoplasm, and (ii) Φ_*d*_(***x***, *t*) = *ϕ*_*h*_(***x***, *t*) − Φ_*e*_(***x***, *t*) which is the difference of the volume fractions of heterochromatin and euchromatin. Note that *ϕ*_*d*_ < (>)0 for the euchromatin (heterochromatin) rich phase, and is therefore analogous to an order parameter. In terms of the chromatin composition variables, *ϕ*_*n*_ and *ϕ*_*d*_, the free energy density at any point ***x*** can be expressed non-dimensionally as (refer SI section S1.4 for details on non-dimensionalization), Note that for a euchromatin (heterochromatin) rich phase *ϕ*_*d*_ < (>)0.

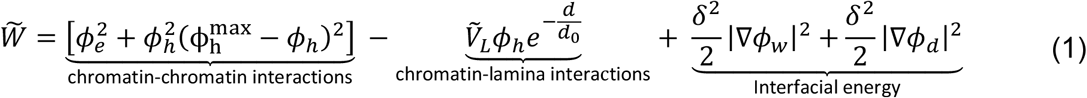

The first term, which is a Flory-Huggins type free energy density for chromatin, defines the competition between the enthalpy of the chromatin-chromatin interactions and entropic contributions of chromatin configuration. This term gives rise to the double-well potential describing the energy landscape of the possible chromatin distribution. The potential surface is visualized in Figure 1b as a contour plot with well locations as *ϕ*_*h*_ = 0 (euchromatin phase) and 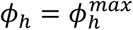 (heterochromatin phase). The well towards the bottom in Figure 1b corresponds to the heterochromatin phase with a low water content and a higher chromatin compaction.

The methylated histone tails in heterochromatin phase can mediate inter-chromatin interactions via chromatin cross-linkers such as HP1*α* [17-19]. Such chromatin crosslinking lowers the enthalpy resulting in a heterochromatin phase well with a densely packed chromatin. On the other hand, the euchromatin well, corresponding to the energy minima with a higher water content is marked with a more acetylated histone tails with a loosely packed chromatin conformation corresponding to a higher entropy.

The second term captures the interactions between the chromatin and the lamina via chromatin anchoring proteins (LAP2*β*, emerin, MAN1, etc.) [20] with parameter 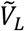 denoting the rescaled strength of these anchoring interactions. Notably, these interactions are most robust at the nuclear periphery (distance from lamina *d* = 0) and vanish exponentially over a length scale *d*_0_. Since the chromatin domains preferentially associating with the nuclear lamina are linked to transcriptional repression and an increased histone methylation [21-24], the chromatin-lamina interactions are captured specifically towards heterochromatin phase. Lastly, the negative sign permits an energetic preference for the peripheral association of heterochromatin. The last term accounts for the interfacial energy which is not accounted in a Flory-Huggins model and penalizes the formation of sharp interfaces between the dissimilar phases (refer Section S1.2 and S1.4 in SI).

### 2.3 Diffusion kinetics of the nucleoplasm

Thus, the energetic considerations dictate that an initial chromatin configuration (red circle in Figure 1b) spontaneously phase-separates into the two energy wells to minimize the total free energy of the system. The driving force pushing the chromatin composition towards the energy wells is a measure of the gradients of the energy landscape and is called the chemical potential. Thus, the chemical potentials are obtained at each point in space by considering changes in energy density for small changes in the local volume fractions (labeled n or d): 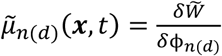. Here, *δ* denotes the functional derivative, or the change in free energy density with respect to the volume fraction. Spatial chemical potential gradients of the nucleoplasm and the epigenetics drive the diffusive flow of the constituents to reduce the overall free energy of the system giving rise to kinetics of the form

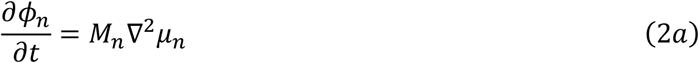

where *M*_*n*_ denotes the mobility of nucleoplasm in the nucleus, which is related to the dissipation that occurs when water flows through the porous nuclear microstructure. An important thing to note here is that nucleoplasm diffusion kinetics in Eq 2a is conservative in nature, i.e. the net amount of water in the nucleus is conserved over time.

### 2.4 Diffusive and reactive kinetics of histone marks

The kinetics of euchromatin and heterochromatin can have two contributions. Phase separation of chromatin creates distinct accumulations of acetylated marked euchromatin and methylated marked heterochromatin. The heterogenous distributions of the epigenetic marks generates diffusive fluxes governed by the gradients in its chemical potential *μ*_*d*_ (first term in Eq 2b). As previously, the diffusion kinetics is conservative in nature, i.e. it does not change the net amount of heterochromatin and euchromatin in the nucleus globally. However, locally, the diffusion kinetics can result in exchange of epigenetic marks – either acetylation or methylation – between neighboring nucleosomes, as shown in the inset in Figure S1. In addition, epigenetic signaling can further result in interconversion of heterochromatin and euchromatin via acetylation or methylation reactions, as shown in Figure S1. Thus, the kinetics of epigenetic regulation and chromatin loop extrusion are incorporated into the dynamics of epigenetic markers, giving rise to a second evolution equation of the form,

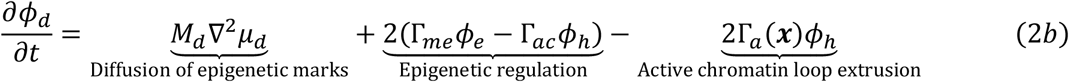

Here, the second term incorporates active first-order reaction kinetics of histone tail acetylation Γ_*ac*_ and methylation Γ_*me*_ leading to interconversion between hetero- and eu-chromatin. The last term accounts for the active chromatin extrusion kinetics and the chromatin state changes resulting from it. As pointed before, while DNA loop extrusion can occur in euchromatin domains, it does not contribute to interconversion of chromatin phases as euchromatin is already transcriptionally active. Contrastingly, at the interface of heterochromatin and euchromatin, DNA loop extrusion can result in activation of otherwise inactive genes. To capture the change in transcriptional status, localized to the hetero-euchromatin interface, we introduce a spatial dependence in the active extrusion rate, Γ_*a*_(***x***), the specific form for which is described in methods (Figure 1b). Important to note here is that the last two terms of equation 2b are responsible for the non-conservative dynamics and can alter the global heterochromatin to euchromatin ratio of the system. More detailed derivation of the chemical potential, contribution of passive diffusion kinetics, epigenetic and active loop extrusion can be found in the methods section.

Having developed the model to capture the spatio-temporal organization of chromatin in the nucleus, we numerically solve Eq 2(a-b) along with the equation defining the chemical potential (Eq S3, SI). The parameters used in the model along with the initial and boundary conditions are described in detail in the SI (Section S8) and listed in Table S1.

## 3 Results

### 3.1 Numerical simulations capture experimentally observed features of chromatin organization

#### In-silico chromatin organization recapitulates cellular observations

We have developed a mathematical model to capture dynamic chromatin organization in the nucleus, in terms of its compaction into the heterochromatic phase or decompaction into the euchromatic phase. We begin by defining the energetics of the chromatin distribution in terms of the entropic-enthalpic balance of chromatin-chromatin interactions, the chromatin-lamina interactions as well as the penalty on the formation of phase boundaries via Eq 1. The gradients in the free-energy landscape (Figure 1b), defined as the chemical potential (refer Eq S3), drive the dynamic evolution of chromatin towards the two energy wells corresponding to the euchromatin and heterochromatin phases via Eq 2a, b. Interconversion of the two phases of chromatin can occur via (a) epigenetic regulation of histone acetylation and methylation, and (b) extrusion of chromatin loops from heterochromatin into euchromatin along the phase boundaries (Eq 2b).

The process of phase separation is initiated by adding a random perturbation to the initially uniform chromatin configuration (as shown in Figure 2a, left panel) which captures the intrinsic intranuclear heterogeneities. As the simulation progresses heterochromatin domains (in red, center panel of Figure 2a) spontaneously nucleate and grow. The evolution ultimately stabilizes resulting in a steady state (right panel of Figure 2a) with a quasi-periodic distribution of stable domains of heterochromatin rich phase 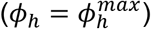 in red and euchromatin rich phase (*ϕ*_*h*_ = 0) in blue. Each of these domains are nearly circular (see Section S2 of SI for a discussion on non-circular lamellar domains) with characteristic sizes. Concomitantly, heterochromatin domains localized to the nuclear lamina (called LADs) of comparable sizes appear in our simulations (Figure 2a).

**Figure 2.**
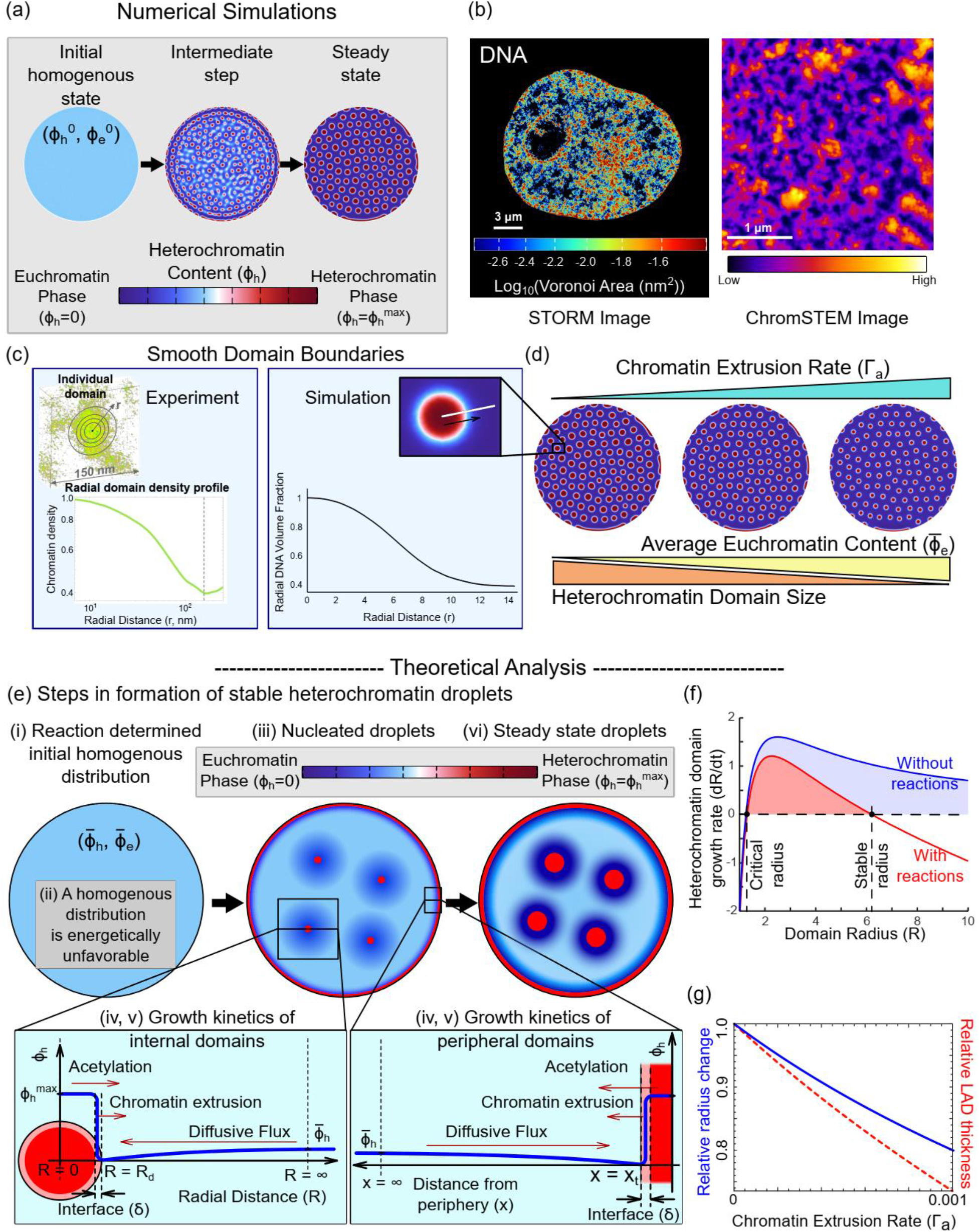
: Numerically predicted chromatin distribution in the nucleus captures the salient features of in-vivo chromatin organization. a) Visualization of the chromatin organization obtained from the simulations. The initial chromatin organization is a homogenous distribution with a small perturbation added, resulting in nucleation of heterochromatin domains (center panel) which grow into heterochromatin domains of characteristic sizes at a steady state. (b) Super-resolution visualizations of chromatin organization observed in-vivo via STORM imaging of HeLa nuclei (left panel, scale bar 3 *μ*m, data previously reported in [14]) and ChromSTEM imaging of BJ fibroblast nuclei (right panel, scale bar 1 *μm*) show that chromatin organization in nucleus is characterized by interspersed heterochromatic domains of comparable sizes. (c) The smooth boundaries of the chromatin packing domains as seen in ChromSTEM observations are captured by the model (d) Numerically predicted trend of sizes of heterochromatin domains as the transcription-mediated chromatin extrusion rate increases. (e) Theoretical analysis of the step-by step events (events ‘i’ through ‘vi’) involved in the nucleation, growth and stabilization of heterochromatin domains at a steady state. (f) Plot of theoretically evaluated growth rate of heterochromatin domains with (red) and without (blue) reactions. Reactions give rise to a stable domain radius. In the absence of reactions, no stable heterochromatin domain length scales are observed. (g) The evaluation of stable radius (blue) and stable LAD thickness (red) as transcription mediated surface reactions are changed.

The meso-scale distribution of chromatin throughout the nucleus predicted by the mathematical model presents a striking qualitative similarity with the experimentally observed distribution of DNA in the nucleus using ChromSTEM, and STORM as reported previously [14] (Figure 2b). Domains of compacted chromatin with a characteristic size are observed via a high histone density distinguished from regions of low histone density (Figure 2b). Lastly, the preferential accumulation of heterochromatin domains along the nuclear periphery seen via STORM imaging (Figure 2b), again with similar size scale, is also in excellent agreement with the experiments.

#### Phase boundaries of heterochromatin domains are smooth

When defining the free energy density of chromatin organization in the nucleus (see Eq S1 in SI), we penalized the formation of sharp interfaces via an interface penalty *η*, defined as the energy cost associated with the formation of the interfaces between heterochromatin and euchromatin phases. As we show in the Section S1.2 and Section S1.4 of the SI, the energy penalty *η* results in the formation of a smooth rather than a sharp interface between the heterochromatin and the euchromatin phases. Numerical simulations of chromatin organization exhibit such smooth interfaces around chromatin domains, as shown in the zoomed in image in Figure 2c (right panel). The width of the interface is controlled by the competition between the interfacial and bulk energy contributions (refer Section S1.4). A smooth description of the chromatin phase boundaries is indeed observed in-vivo (SI Section S1.9). ChromSTEM was used for a 3D characterization of chromatin density around individual heterochromatin domains in a BJ fibroblast nucleus (Figure 2c, left panel). We estimated the average chromatin density within concentric circles emerging from the center of individual domains to the periphery (Figure 2c, Figure S3). The chromatin density was highest at the core of the domain and dropped slowly from the center of the domain to the periphery until the surrounding domains intersect. The smooth decrease in radial density indicates that the chromatin domain boundaries are not abrupt (Figure 2c), in agreement with the numerical simulations.

#### Heterochromatin domain length-scales are determined by reaction kinetics

We next investigate how the size scaling of the heterochromatin domains is regulated by the epigenetic reactions – acetylation and methylation of histones – and transcriptionally active loop extrusion which together can lead to interconversion between heterochromatin and euchromatin. First, we see that in the absence of the epigenetic and active extrusion reactions multiple domains of a characteristic size are not obtained as shown in Figure S7 (detailed discussion in SI Section S5). In this case, although nucleation of multiple heterochromatin domains occurs even without reactions, all of them merge into a single large cluster driven by Ostwald ripening so as to minimize the interface formation.

The model also predicts that the size of the heterochromatin domains in the interior and periphery can be regulated by the epigenetic reaction rates of acetylation and methylation as shown in Figure S4 (SI Section S2). We see that as methylation increases the size of the interior domains increases too. On the other hand, increase in acetylation results in the formation of smaller heterochromatin domains. The trends followed by the domains towards the interior of the nucleus are replicated by the LADs as well. Lastly, we identify that the size scales of the domains – the domain radii in the interior of the nucleus and the LAD thickness along its periphery – depend on the level of transcription governed chromatin extrusion rate Γ_*a*_ (Figure 2c). We note that, as the transcription (Γ_*a*_) is increased, the sizes of the heterochromatin domains decrease, both in the interior as well as at the periphery. At the same time, we also note that as chromatin extrusion rate is increased, the average volume fraction of heterochromatin 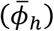 in the nucleus decreases, while that of euchromatin 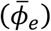 increases.

### 3.2 Theoretical analysis predicts how the heterochromatic domain and LAD sizes depend on epigenetic and transcriptional regulation

Next, we theoretically predict an explicit dependence of the sizes of interior heterochromatic domains and LADs on epigenetic and transcription reactions and the diffusion kinetics of the epigenetic marks.

#### Determination of the average eu- and heterochromatin volume fractions

Intuitively, in the presence of more repressive methylation the overall heterochromatin content in the nucleus should increase, while in higher histone acetylation conditions the overall euchromatin content will increase. Thus, the epigenetic reactions independently determine the average volume fractions of each form of chromatin, thereby breaking the detailed balance condition where the free energies of each phase determine their relative abundance in a thermodynamic equilibrium. A mathematical relation between the average volume fraction of each chromatin phase and the epigenetic reaction parameters can be determined by averaging the chromatin evolution equation (Eq 2b) at a steady state (i.e. 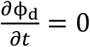). In the absence of transcription driven chromatin extrusion (i.e. 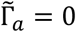), we see that the epigenetic kinetics regulates the average heterochromatin content of the nucleus as, 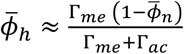 (Eq. S11, refer SI Section S3 for more details).

The presence of transcription-driven loop extrusion kinetics (i.e., 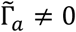 in Eq 2b) further augments the deviation from thermodynamic equilibrium (i.e., the breaking of detail balance) via surface reactions that actively extrude DNA at the interface of heterochromatic domains. In the presence of transcription, the average heterochromatin (and euchromatin) content in the nucleus becomes,

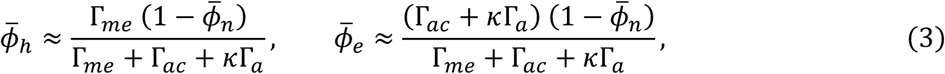

where *κ* is a function of 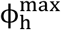, volume fraction change across the interface Δ*ϕ*, and the length of the interface between the two chromatin phases (refer SI Section S3 for derivation). Since DNA loop extrusion converts heterochromatin into transcriptionally active euchromatin form, as extrusion rate Γ_*a*_ increases, the average heterochromatin content decreases. Thus, the epigenetic regulation along with transcription determine the mean chromatin composition of the nucleus given as 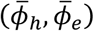 as shown in Figure 1b (light blue circle).

#### Biophysical origin of characteristic heterochromatin domain size

Next, we show that the average composition of the two chromatin phases, shown in Figure 2d(i), plays a key role in the emergence of the characteristic sizes of the heterochromatin domains. To illustrate this, we first observe that the mean chromatin composition 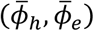 lies in neither of the energy wells as shown in Figure 1b (light blue circle) and is thus energetically unfavorable. The need to reduce the total free energy in the nucleus drives the system to phase separate by nucleating heterochromatin domains (Figure 2d(iii)) corresponding to the red energy well labeled heterochromatin in Figure 1b surrounded by euchromatin domains corresponding to the dark blue energy well labeled euchromatin. The events entailing the individual steps in the nucleation and growth of a single droplet of heterochromatin due to phase separation, as shown in Figure 2d, are as follows:

1. Due to phase separation, the heterochromatin volume fraction immediately outside the droplet is *ϕ*_*h*_ = 0 corresponding to the euchromatic energy well. Far away from the droplet, the mean composition 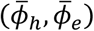 remains undisturbed. The resulting spatial gradient in the chromatin composition (blue curve in Figure 2d(iv)) sets up a diffusive flux of heterochromatin into the droplet, allowing it to grow.
2. On the other hand, within the heterochromatin droplet (with 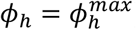) histone acetylation reactions will allow conversion of heterochromatin inside the droplet into euchromatin outside. Active chromatin loop extrusion further adds to the heterochromatin outflux. Together loop extrusion and acetylation oppose the diffusive influx of heterochromatin and thereby reduce the size of the droplet (Figure 2d).
3. Based on the above observations, the rate at which the nucleated heterochromatin droplet grows can be written in terms of the balance of reaction-diffusion gradient driven influx and acetylation and transcription driven outflux of heterochromatin as (refer SI Section S4, Eq S13),

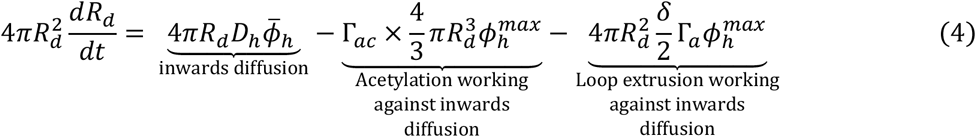

where *δ* is the width of the interface, which is in turn related to the length scale obtained via the competition between the interfacial energy and chromatin-chromatin interaction (refer methods, Eq 1), and *D*_*h*_ is the diffusivity of heterochromatin, which is determined by the mobilities of nucleoplasm *M*_*n*_ and epigenetic marks *M*_*d*_. The resulting evolution of the droplet growth rate (*dR*_*d*_/*dt*) as the radius of the droplet increases is shown in Figure 2e. Notice the two fixed points (Figure 2e, labelled critical and stable radius) where *dR*_*d*_/*dt* = 0. Beyond the critical radius the domains grow in size.
4. The second fixed point (stable radius) corresponds to the rescaled steady state (i.e., *dR*_*d*_/*dt* = 0) heterochromatin domain size as determined by the active epigenetic and the transcriptional regulation in tandem with passive diffusion, and can be written as (derivation shown in SI, Section S4, Eq S18),

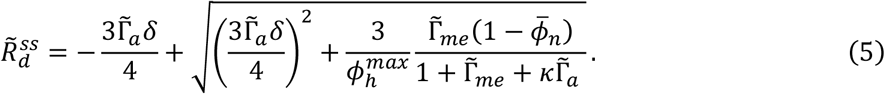

From Eq 5, we observe that the steady state droplet radius 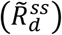 depends on both diffusion and reaction kinetics. With increase in methylation, 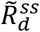 increases implying bigger heterochromatin domains. On the other hand, with increase in either the acetylation or transcription-mediated loop extrusion the steady state radius decreases. The quantitative dependence of the steady state radius on transcriptional kinetics is shown in Figure 2f (blue solid line). Thus, our theory predicts an increase in the sizes of compacted chromatin domains in the interior of the nucleus upon inhibition of transcription.

#### Regulation of LAD thickness by epigenetic reactions and transcription

The size dependence of chromatin domains along the nuclear periphery can be similarly determined by the balance of reaction, transcription, and diffusion kinetics for the LADs. The affinity of chromatin to the nuclear periphery due to the chromatin-lamina interactions in Eq 1 induces a preferential nucleation of LADs. A schematic representation of heterochromatin compaction along the nuclear periphery resulting in LAD growth is shown in Figure 2d. As with the interior heterochromatin droplet, phase-separation drives the heterochromatin compaction 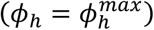 within the LADs, while the chromatin immediately outside corresponds to the euchromatin energy minimal well (*ϕ*_*h*_ = 0). Far away from the peripheral LAD nucleation sites, the chromatin composition remains undisturbed at the average composition of 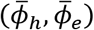. The variation of chromatin composition with distance from nuclear periphery is shown in Figure 2d (blue line). Like in the case of the interior heterochromatin droplets, the heterochromatin composition gradient driven diffusive influx is balanced by the epigenetic and transcriptional regulated heterochromatin outflux, which determines the rescaled steady-state thickness of the LADs (refer to the SI, Section S7, Eq. S20),

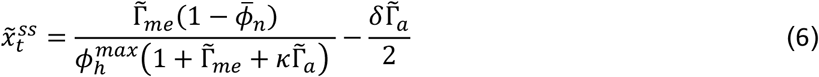

As with the interior domains, we observe that the LADs become thicker with increase in methylation, while they become thinner with increasing acetylation or chromatin extrusion rates. A quantitative dependence of steady state LAD thickness on transcription rate based on Eq 6 is plotted in Figure 2f (red dashed line). Our theory predicts an increase in the sizes of LADs along the nuclear periphery upon inhibition of transcription. While the theoretical analysis helps develop a fundamental biophysical understanding of the role of energetics and kinetics in chromatin phase separation, a nucleus-wide chromatin organization and its dynamic evolution can only be obtained numerically.

### 3.3 Loss of transcription results in increase in heterochromatin domain size and LAD thickness

Next, we use the in-silico model to make testable quantitative predictions of the meso-scale chromatin organization in the nucleus. We also report the in-vivo nuclear chromatin reorganization upon transcription inhibition using complimentary STORM [14] and ChromSTEM – on nuclei from multiple cell lines. The choice of the parameters for rates of acetylation 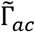, methylation 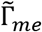, and the strength of chromatin-lamina interactions 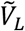, were held constant for all the following simulations, and the choice of the level of spatial noise is discussed in the SI Section S8. We calibrate the active chromatin loop extrusion rate Γ_*a*_ to obtain an *in-silico* change in the interior domain sizes quantitatively comparable to that observed upon transcriptional inhibition. The calibrated model is then used to predict the change in LAD thickness due to inhibition of transcription, which upon comparison with experimental images serves to validate the model. A schematic for the workflow utilized to calibrate and cross-validate the model predictions in the interior and along periphery of the nucleus is shown in Figure S11.

#### Reduction in size of heterochromatin domains upon transcriptional inhibition is observed in-vivo over multiple cell-lines

ChromSTEM was used to obtain super-resolution images in terms of statistical descriptions of chromatin packing domains for BJ fibroblasts. ChromSTEM allows the quantification of 3D chromatin conformation with high resolution [25]. ChromSTEM mass density tomograms were collected for BJ fibroblasts treated with Actinomycin D (ActD) (Figure 3a, center panel) and compared to DMSO treated mock controls (Figure 3a, left panel) to evaluate the average size and density of chromatin packing domains. We have previously demonstrated that chromatin forms spatially well-defined higher-order packing domains and that, within these domains, chromatin exhibits a polymeric power-law scaling behavior with radially decreasing mass density moving outwards from the center of the domain [26]. As the ChromSTEM intensity in the reconstructed tomogram is proportional to the chromatin mass density, we estimated the size of the domains based on where the chromatin mass scaling and the radial chromatin density deviate from their predicted behavior (discussed in methods, Section 2.5). Based on the statistical analysis of individual packing domains, in a single tomograph shown in Figure 3a, we observed 71 domains in DMSO and 48 domains in the ActD-treated nucleus. Of the identified domains, the average domain radius (± S.E) of BJ cells treated with DMSO and ActD was estimated to be 103.5 ± 4.73 nm and 129.7 ± 6.78 nm, respectively (Figure 3a, right panel), representing a 20.2% increase in size. Overall, fewer domain centers, and larger chromatin packing domains were experimentally observed upon ActD treatment compared to the control.

**Figure 3.**
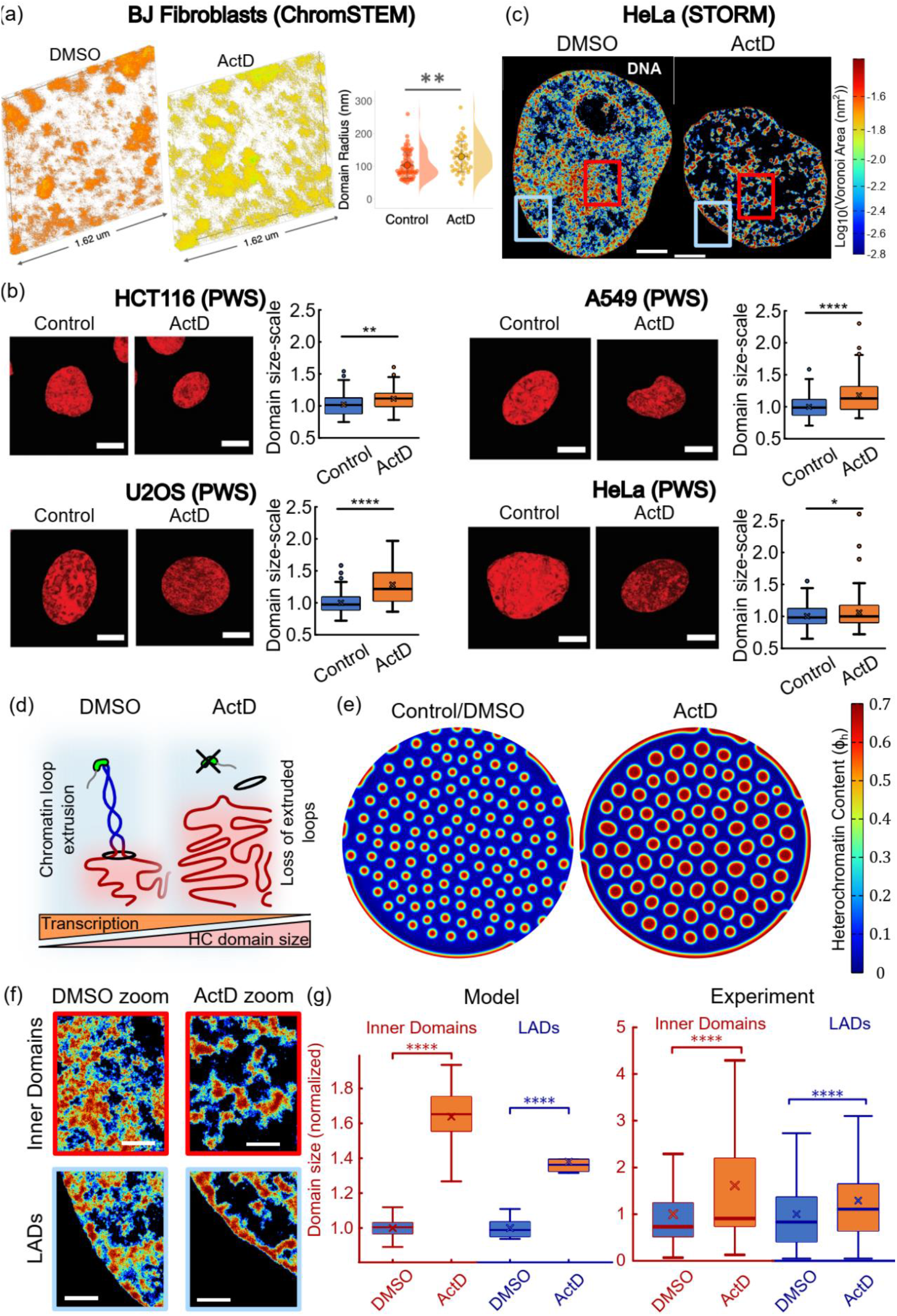
: Heterochromatin domains increase in size after transcription inhibition. (a) ChromSTEM tomogram reconstructions for DMSO (left panel, chromatin density in orange) and ActD treated (center panel, chromatin density in yellow) BJ Fibroblast cells. The distribution of the radius of packing domains, for BJ cells treated ActD (right panel, n=48 domains) show a significant increase in size compared to control (n = 71 domains, ∼1.25 times, p <0.01). (b) Representative live-cell PWS images for four cell lines demonstrate significant reorganization of chromatin within 1 hr of treatment with ActD. Scale bars represent 5 μm. Box plots show the comparison of domain sizes between DMSO control and ActD treated cells. (n > 100 cells/ condition for each cell type; *p ≤ 0.05, **p≤0.005, ****p ≤ 0.0001) (c) Heatmap density rendering of super-resolution images of DNA in DMSO control (left panel) and ActD (right panel) treated HeLa nuclei. All scale bars - 3*μ*m. (d) A schematic representation of loss of chromatin loop extrusion due to absence of RNAP2. The size of the heterochromatin domains (red region) increases. (e) Numerical prediction of distribution of heterochromatin domains (in red) in a nucleus with DMSO control and without transcription mediated chromatin extrusion (ActD). (f) Zoomed in views of DMSO and ActD treated nuclei localized to the nucleus interior (top panels) and the periphery (bottom panels). Red and blue boxes shown in (c) are zoomed into. All scale bars -1*μ*m. (g) Boxplots showing the distribution of radii of heterochromatin domains predicted numerically. ActD nuclei have a mean interior domain radius 1.63 times that of DMSO nuclei (unpaired two tail t-test, p = 0). ActD nuclei also have a mean LAD thickness 1.37 times that of DMSO nuclei (unpaired two tail t-test, p = 0). Right panel shows the quantification of the distribution of radii of heterochromatin domains in DMSO and ActD nuclei (*n* ∼ 10^3^ loci in 19 nuclei for DMSO treatment and 20 nuclei for ActD treatment). ActD treated nuclei show a significantly greater heterochromatin domain radius (∼1.61 times, unpaired two tail t-test, p = 0). Quantification of the distribution of LAD thickness in DMSO and ActD nuclei. ActD treated nuclei show a significantly greater LAD thickness (∼1.3 times, unpaired two tail t-test, p = 0.006).

In addition to evaluating domain properties using ChromSTEM, we utilized live-cell partial wave spectroscopy (PWS) imaging to observe the change in chromatin organization after transcription inhibition in various cell lines (Figure 3b). The PWS images demonstrate a significant reduction in average chromatin packing scaling upon ActD treatment in live cells across four different cell types. Next, the size of the domains is quantitatively approximated via polymer scaling relationships discussed in Section S1.12 (Supplementary Information) [25, 27]. The quantification of the domain sizes (boxplots in Fig 3b) shows that, for all cell types studied, packing domains are larger for upon transcription inhibition with ActD treatment – in agreement with the ChromSTEM results on BJ fibroblasts.

Additionally, we have previously used STORM imaging to observe the nucleus wide changes in chromatin organization caused by transcription abrogation in HeLa nuclei after ActD treatment [14]. Heatmaps of chromatin density obtained via Voronoi tessellation-based color-coding of STORM images (see [14] for analysis) are shown in Figure 3c. The zoomed in images of heatmaps of the chromatin cluster density (Figure 3f) clearly show the increasing heterochromatin domain sizes when RNAPII activity is inhibited, in agreement with our theoretical and numerical predictions (Figure 2c, d). Importantly, we see that the changes in chromatin organization occur not only in the interior domains of the nucleus but also along its periphery (Figure 3f, g).

Altogether these complementary imaging techniques establish that nucleus wide increase in sizes of compacted chromatin domains occurs upon the loss of transcription in a wide range of cell lines.

#### Analysis of the change in size of interior heterochromatin domains provides interfacial loop extrusion rates

The chromatin cluster density maps obtained above were further analyzed to quantify the sizes of heterochromatin domains after DMSO and ActD treatment. A density-based threshold was used to isolate the high-density heterochromatin regions, which were then clustered via a density based spatial clustering algorithm (see SI Section S1.6) and further sub-classified into LADs and interior domains depending on the distance from nuclear periphery. The quantitatively extracted distribution of interior heterochromatin domain radii for DMSO and ActD treated nuclei (see SI Section S1.7) shows that their mean radius after transcription inhibition was nearly 1.61 times that in DMSO controls (Figure 3g)).

Indeed, our model (Section 3.1, Figure 2c) predicts that loss of transcription results in increased heterochromatin domain size. This is because under control conditions, extrusion of heterochromatin phase into euchromatin occurs. We assume, based on previous experimental findings [14], that the presence of RNAPII activity drives the supercoiling of the DNA loop, thereby extruding it from the heterochromatin phase into the euchromatin phase at the phase boundaries (Figure 3c, left panel). However, when RNAPII is inhibited with ActD treatment (Figure 3c, right panel), the absence of this driving force for loop extrusion keeps more DNA in the heterochromatin phase thereby increasing the domain sizes. The *in-silico* chromatin distribution predicted under control (left panel) and transcription inhibited (Γ_*a*_ = 0, right panel) conditions is shown in Figure 3a. The phase separated heterochromatin domains 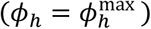 are shown in red in a loosely compacted euchromatin background (blue, *ϕ*_*h*_ = 0). We quantify the change in the sizes of the heterochromatin domains predicted by the model as the active extrusion rate Γ_*a*_ is parametrically varied. The value of Γ_*a*_ under control conditions is chosen (Table S1) such that the change in the interior domain sizes with respect to transcription inhibition (with Γ_*a*_ = 0) is quantitatively the same as observed experimentally.

#### The model predicts changes in LAD thickness due to transcriptional inhibition with no additional parameters

Next, we quantitatively validate the choice of Γ_*a*_ under control conditions by comparing the predicted change in LAD thickness against that quantified from the STORM images. Our theoretical predictions discussed in Section 2.1 (Eq 6) show that the reduction in transcription increases the thickness of the LADs reflecting the behavior predicted in the interior of the nucleus (Figures 2d and 2g). Our simulations of chromatin distribution in the nucleus (Figure 3e) show that inhibition of transcription (Γ_*a*_ = 0) results in thicker LADs. Of note, the chromatin-lamina interaction strength (*V*_*L*_) stays unchanged between the two simulations. Yet, we see a higher association of chromatin with the periphery. Upon quantitative comparison (Figure 3b, right panel) we see that the LADs grow approximately 1.37 times thicker upon loss of transcription.

To validate this prediction, we compare the predicted change in LAD thickness with that quantified from *in-vivo* STORM imaging. (Figures 3g, refer to methods section for procedure). The quantified comparison of LAD thickness between DMSO and ActD nuclei (Figure 3g) shows nearly 1.3 times increase upon ActD treatment, in close quantitative agreement with the model prediction. Overall, with both model predictions and cellular observations, our results suggest that impairment of transcription plays a significant role in determining the size scaling of the interior heterochromatin domains and LADs.

### 3.4 Transcription inhibition results in movement of DNA from the euchromatic into heterochromatic regions

We next enquire how, in addition to altering the size of the compacted domains, abrogation of transcription changes the extent of DNA packing. For this we analyzed the chromatin distribution in HeLa nuclei under DMSO and ActD treatments from STORM images previously generated [14]. Under control conditions the distribution of DNA is qualitatively more homogenous while ActD treated nuclei exhibit more isolated distinct domains of compacted chromatin surrounded by region of very low chromatin density (Figure 4a). For quantification, we plot the chromatin intensity along a horizontal line chosen to run across two heterochromatin domains with euchromatin between them (see zoomed images in Figure 4b, blue horizontal line). The chromatin intensity, plotted in Figure 4c (in blue) shows that even in the euchromatin region, the DNA presence is substantial. On the other hand, chromatin intensity across a horizontal line chosen across a heterochromatin domain in ActD nucleus shows much steeper gradient outside the domain.

**Figure 4.**
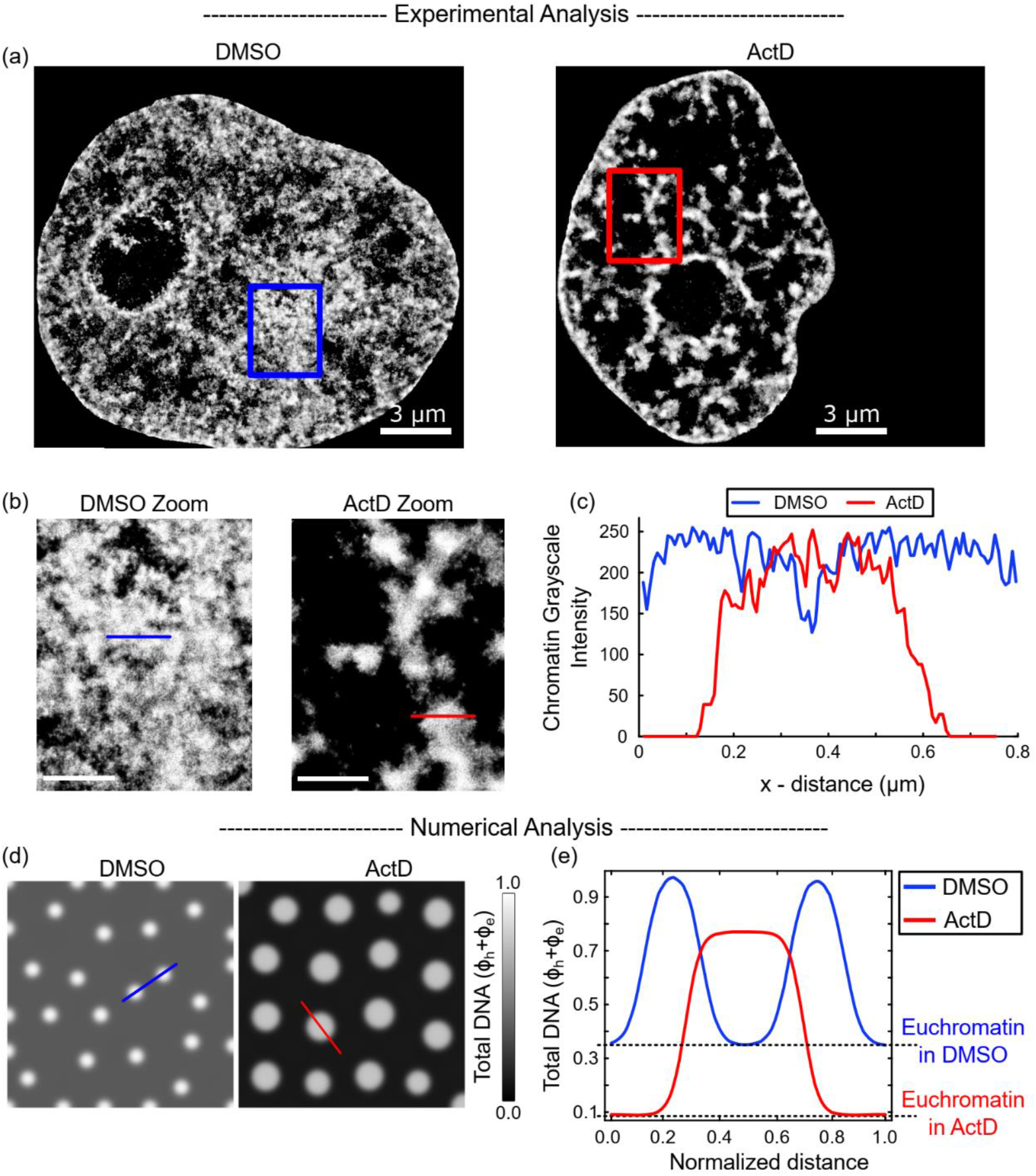
: Loss of transcription reduces the amount of DNA in euchromatic phase. (a) Grayscale heatmap density rendering of super-resolution images of DNA in control (DMSO, left panel) and actinomycin D (ActD) treated HeLa nuclei. All scale bars - 3*μ*m. (d) Zoomed in views of DMSO and ActD treated nuclei. Boxes shown in (a) are zoomed into. All scale bars - 1*μ*m. (e) Along the blue (DMSO) and red (ActD) line segments, we plot the chromatin heatmap intensity (corresponding to the total DNA content) for the DMSO-treated control nucleus (in blue) and ActD-treated nucleus (in red). The DMSO-treated nucleus shows a wider distribution of small heterochromatin domains, while the ActD treated nucleus shows a greater compaction with isolated large heterochromatin domains. (a) Numerical prediction of distribution of total DNA (in grayscale) in a nucleus with (DMSO) and without (ActD) transcription mediated chromatin extrusion. (b) Distribution of total DNA content along the blue (red) line in (a) under DMSO (ActD) treatment.

The increased presence of DNA in the euchromatic phase observed experimentally is captured by the simulations. The distribution of DNA (measured as the sum of volume fractions of the chromatin phases, *ϕ*_*e*_ + *ϕ*_*h*_) in a nuclear region far from LADs is plotted in Figure 4d for control and transcription inhibited in-silico nuclei. We see that the euchromatic phase (outside white circles) is darker when transcription is inhibited, indicating the presence of much lesser DNA than in control euchromatin. A quantification of the total DNA along cut-lines chosen in the control and ActD in-silico nuclei confirm the observations (Figure 4e).

Since the lack of transcription inhibits chromatin loop extrusion from heterochromatin into euchromatin, we see a reduced density of DNA in the euchromatin phase of the nucleus under ActD conditions. Further, due to the lack of chromatin extrusion out of the heterochromatin domains when transcription is inhibited, we also observe that they are larger in size. Thus, transcription, via chromatin loop extrusion, results in removal of DNA from compacted heterochromatin region by converting it into active euchromatin form.

Taken together, our results suggest that transcription not only affects the scaling of the lengths (radius or thickness) of the heterochromatin domains, but also significantly changes the relative amounts of DNA in the euchromatin and heterochromatin phases.

### 3.5 Excessive chromatin loop extrusion reduces the sizes of chromatin domains

We have established that change in transcription activity affects the global chromatin organization of the nucleus via altered loop extrusion. In turn, chromatin loop extrusion is initiated by the loading of cohesin onto DNA via a balance between cohesin loaders such as NIPBL and cohesin unloaders like WAPL (Figure 1c [2-4, 28]). If the chromatin loop extrusion is responsible for the global chromatin reorganization, altering the cohesin loading/unloading balance must also result in chromatin reorganization. Thus, next, we study the chromatin arrangement in WAPL-deficient (WAPLΔ) nuclei marked by increased levels of loaded cohesin.

#### Decrease in domain sizes due to WAPL deficiency quantitatively predicts excessive chromatin loop extrusion

*In vivo*, WAPL depletion causes an accumulation of large amounts of cohesin on chromatin [29]. This results in a much more homogenous distribution of DNA, which was previously termed “blending” due to excessive extrusion of chromatin loops, as shown schematically in Figure 5a [14]. In our mathematical model, WAPL deficiency is simulated as an increase in the rate of chromatin extrusion (Γ_*a*_). Based on the theoretical size scaling of the interior heterochromatin domains and LADs, as seen from Eq. 5 and Figure 2f, our model predicts that increase in Γ_*a*_ would result in a decrease in the radius of the steady state heterochromatin domains (Figure 5b).

**Figure 5.**
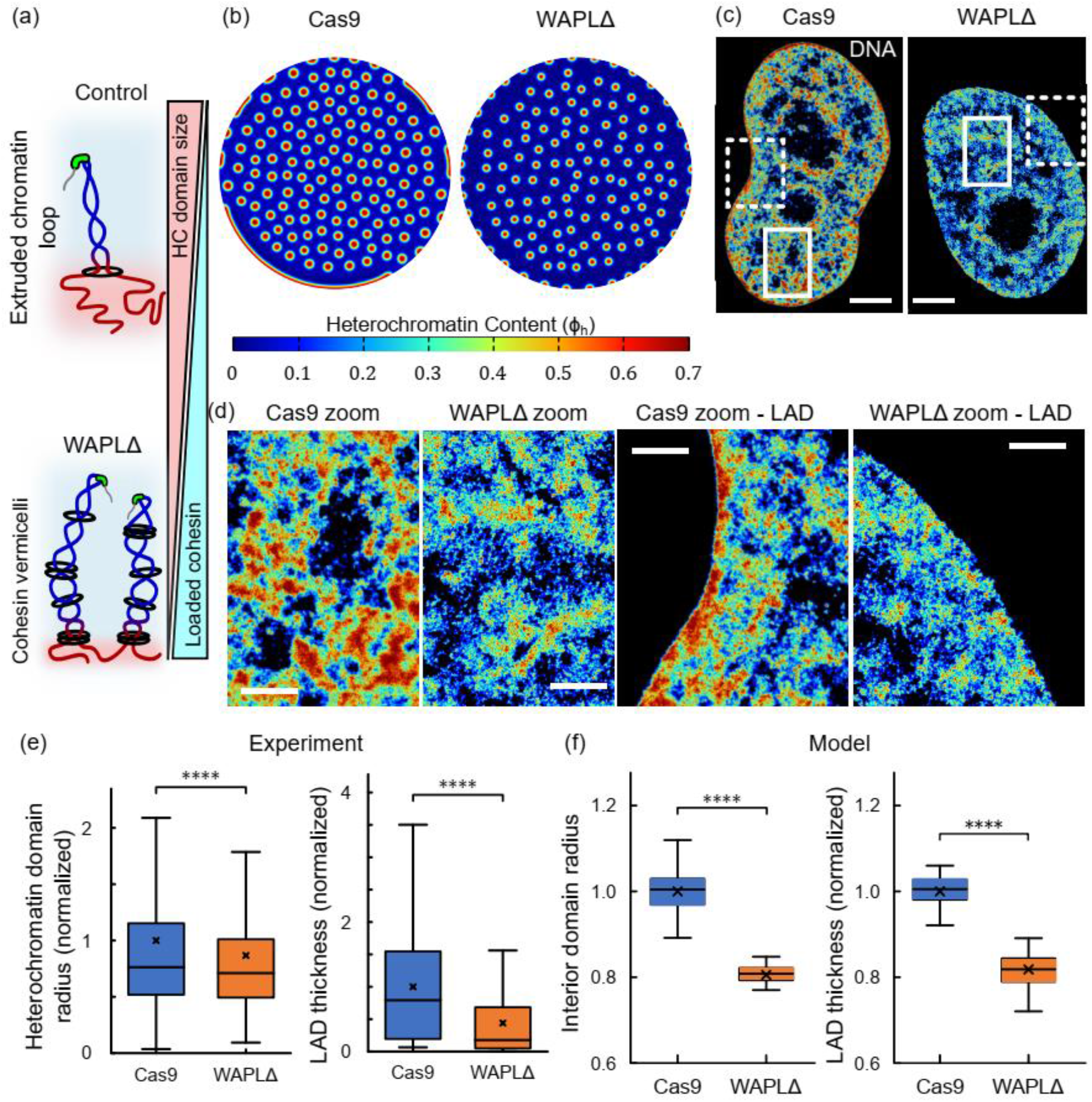
: Heterochromatin domains become smaller upon WAPL-depletion. (a) A schematic representation of the formation of chromatin loops. After WAPL-depletion there is an increased loading of cohesin resulting in excessive chromatin loop extrusion (b) Numerical prediction of distribution of heterochromatin domains in the interior and the LADs along the periphery (all domains in red) in a nucleus without (Cas9) and with (WAPLΔ) cohesin unloading disruption. (c) Heatmap density rendering of super-resolution images of DNA in (d) control (Cas9, left panel) and WAPL knock-out (WAPLΔ) treated HeLa nuclei. All scale bars - 3*μ*m. (d) Left panel shows zoomed in views of Cas9 and WAPLΔ treated nuclei focusing on the heterochromatin domains in the interior of the nucleus. White solid box shown in (c) is zoomed into. All scale bars - 1*μ*m. Right panel shows zoomed in views of Cas9 and WAPLΔ treated nuclei focusing on the LAD domains along the nuclear periphery. White dashed box shown in (c) is zoomed into. All scale bars - 1*μ*m. (e) Quantification of heterochromatin domain radius in the interior of Cas9- and WAPLΔ-treated nuclei. (*n* ∼ 10^3^ loci in 6 nuclei for Cas9-treatment and 7 nuclei for WAPLΔ treatment). WAPLΔ treated nuclei exhibit a significantly lower (∼ 0.86 times) mean heterochromatin radius (unpaired two-tailed t-test, p ∼ 1e-10). Quantification of LAD thickness along the periphery of Cas9- and WAPLΔ-treated nuclei. (*n* ∼ 10^2^ loci in 6 nuclei for Cas9-treatment and 7 nuclei for WAPLΔ treatment). WAPLΔ treated nuclei exhibit a significantly lower (∼ 0.43 times) mean LAD thickness (unpaired two-tailed t-test, p ∼ 1e-12). (f) Boxplot in left panel shows the distribution of domain radii predicted numerically. WAPLΔ nuclei have a mean domain radius 0.8 times that of Cas9-treated nuclei (unpaired two-tailed t-test, p = 0). Boxplot in right panels shows the distribution of LAD thicknesses predicted numerically. WAPLΔ nuclei have a mean LAD thickness 0.82 times that of Cas9-treated nuclei.

STORM images of HeLa nuclei without (labeled Cas9) and with WAPL-deficiency previously revealed genome-wide changes in the chromatin organization induced by excessive loading of cohesin (Figures 5c and 5d) [14]. A visual comparison between representative zoomed-in regions (white boxes in Figure 5c) demonstrates the reduction of heterochromatin domain sizes in the interior of the nuclei in WAPLΔ nuclei (Figure 5d). Using clustering analysis (refer Section S1.6 and S1.7), we quantify the altered chromatin domain sizes in control and WAPLΔ HeLa cell nuclei. We observe that WAPLΔ nuclei with increased chromatin blending have heterochromatin domains with a mean radius approximately 15% smaller than control nuclei (Figure 5g).

*In-silico*, we parametrically vary the active chromatin extrusion rate Γ_*a*_ above the control level (determined in Section 3.3). The value of Γ_*a*_ for WAPLΔ nuclei is chosen (Table S1) such that the decrease in the size of interior heterochromatin domains reduces by 15% to agree with the experimental observation (Figure 5f).

#### Model predicts the loss of LADs due to WAPL-deficiency with no additional parameters

As discussed previously (Figure 2f), the model predicts that the effects of chromatin extrusion observed in the interior domains of the nucleus are replicated along the nuclear periphery. Simulation of nuclear chromatin organization (Figure 5b) reveals that by changing only the rate of chromatin extrusion Γ_*a*_, keeping all other parameters including chromatin-lamina interaction potential *V*_*L*_ constant, we see a reduction in the association of chromatin with the lamina. Specifically, a 2.5-fold increase in Γ_*a*_ calibrated to occur due to WAPL-deficiency predicts a 51.2% decrease in the average LAD thickness, as shown in Figure 5c.

The predicted change in LAD thickness is consistent with previous experimental observations and was further quantitatively validated by measuring the thickness of LADs in STORM images of control and WAPLΔ nuclei (Figure 5e) [14]. A reduction in the sizes of domains, as seen in the nucleus interior, can also be observed at the nuclear periphery, as shown in a representative zoomed in region (white dashed boxes in Figure 5c) in Figure 5d. The mean thickness of the LADs at the nuclear periphery is approximately 57% smaller for WAPLΔ nuclei (Figure 5h) as compared to the control-treated nuclei – in close agreement with the model predictions.

Together, these results confirm that the meso-scale spatial chromatin organization is strongly regulated by the chromatin loop formation, and this effect can be modulated not only by the transcription activity, but also by altering the extent of loading or unloading of cohesin rings on the DNA. These results provide further evidence for the link between transcriptional regulation and nucleus-wide chromatin distribution via chromatin loop extrusion.

### 3.6 Chromatin blending in WAPL deficient cells is blocked by transcription inhibition

Since we have established, via both quantitative analysis of experimental data and simulations, that extrusion of chromatin loops is governed by both cohesin loading/unloading balance and RNAPII mediated transcription, a question of their tandem role emerges.

#### Individual steps in chromatin loop extrusion

To simulate the individual effects of cohesin loading and transcriptional activity, we decompose the overall active chromatin extrusion rate into its distinct constitutive steps. The individual steps involved in the process of chromatin loop extrusion from heterochromatin into euchromatin (as discussed previously in Section 1) are shown in Figure 6a. As a first step, a balance between the loading of cohesin via NIPBL/MAU2 [28] on chromatin occurring at a rate Γ_*l*_ and its unloading via by WAPL/PDS5 [2-4] occurring at a rate Γ_*ul*_ results in the association of cohesin rings with chromatin at an overall rate Γ_*coh*_ = Γ_*l*_ − Γ_*ul*_. In other words, Γ_*coh*_ denotes the overall rate of cohesin loading on DNA. The entrapment of DNA by cohesin is followed by the extrusion of supercoiled loops of chromatin via DNA supercoiling by the RNAPII mediated transcription, at a rate denoted by Γ_*tr*_. Thus, as shown in Figure 6a, by assuming a first-order reaction kinetics for both steps, the overall rate of active chromatin extrusion Γ_*a*_ at the interface of heterochromatin and euchromatin is proposed to be multiplicatively decomposed as,

**Figure 6.**
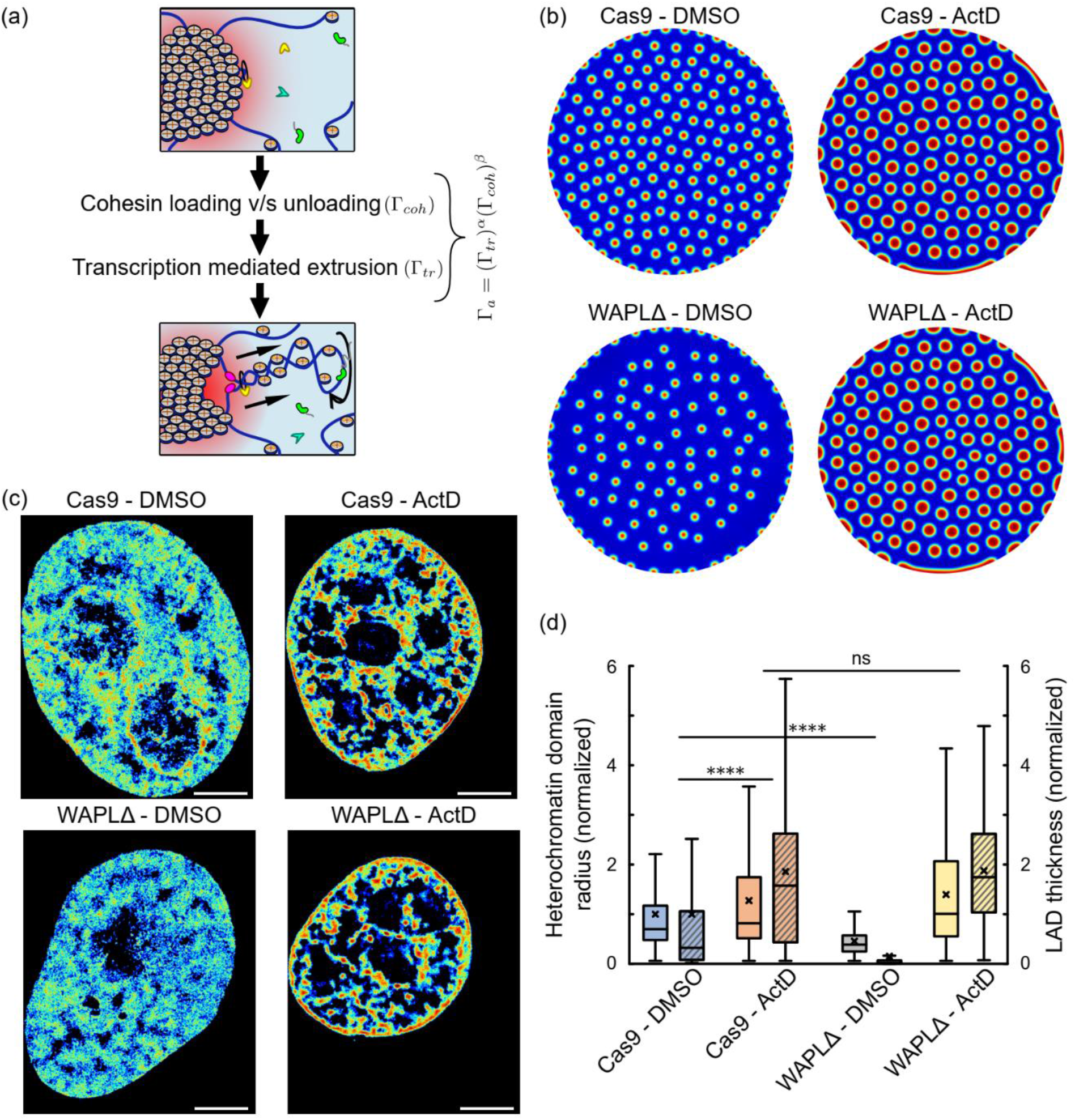
: Simultaneous roles of transcription inhibition and cohesin imbalance (via disabling cohesin unloading WALPΔ) are studied. (a) Schematic showing the associative sub-steps of chromatin extrusion incorporating cohesin loading v/s unloading balance and active transcriptional work done by RNAPII. The rate of active extrusion of chromatin loops (Γ_*a*_) is determined by both sub-steps. (b) Numerical prediction of distribution of heterochromatin domains in the interior and the LADs along the periphery (all domains in red) in a nucleus in control (Cas9-DMSO treatment, top-left panel), transcription inhibited (Cas9-ActD, top right), WAPL knock-out treated (WAPLΔ-DMSO, bottom left) and simultaneous WAPL knock-out along with transcription inhibition treated (WAPLΔ-ActD, bottom right). (c) Heatmap density rendering of super-resolution images of DNA in control (Cas9-DMSO treatment, left panel), transcription inhibited (Cas9-ActD, center left), WAPL knock-out treated (WAPLΔ-DMSO, center right) and simultaneous WAPL knock-out along with transcription inhibition treated (WAPLΔ-ActD) HeLa nuclei. All scale bars - 3*μ*m. (d) Quantification of heterochromatin domain radius in the interior (plain colored boxes) as well as the LAD thickness along the nuclear periphery (hatched boxes) of Cas9-DMSO, Cas9-ActD, WAPLΔ-DMSO and WAPLΔ-ActD treated nuclei (*n* ∼ 10^3^ loci in 12 - 15 nuclei for each treatment). As previously, ActD treated nuclei exhibited a significantly increased domain size (unpaired two-tailed t-test, p ∼ 1e-10) while WAPLΔ treated nuclei exhibit a significantly lower mean heterochromatin radius (unpaired tw-tailed t-test, p ∼ 1e-10). However, the differences between Cas9-ActD treated and WAPLΔ-ActD treated nuclei was insignificant (unpaired two-tailed t-test, p ∼ 0.9).

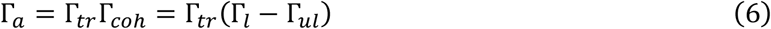

In addition to the extrusion of loops via RNAPII mediated DNA supercoiling activity [4, 8, 14, 30-32], *in vitro* experiments proposed that cohesin once transiently loaded onto DNA, could independently drive the formation of loops via its ATPase machinery [5, 7, 33-35]. Cell based experiments demonstrated that in WAPLΔ cells, clusters of cohesin in WAPLΔ cells assemble together into vermicelli-like structures and these structures disappear upon transcription inhibition, but not upon partial loss of cohesin [14]. These results, taken together, present strong evidence for the important role of transcription in powering cohesin mediated loop extrusion. While the relative role of cohesin’s motor activity and transcription in loop extrusion inside cells remains to be determined, here we focus on the latter given the previous in vivo experimental findings. We indeed show that a kinetic model captured by Eq. 6 sufficiently explains the effect of extrusion of the specific chromatin loops extending from transcriptionally silenced heterochromatin into genetically active euchromatin on determining the meso-scale chromatin domain sizes.

#### Computational model predicts the tandem roles of transcription and cohesin loading

The chromatin organization is simulated in a nucleus under control and transcription inhibition treatments for nuclei with and without WAPL deficiency. The chromatin organization in a control nucleus (labelled Cas9-DMSO), simulated via parameters listed in Table S1 is shown in Figure 6b, top-left panel. The individual inhibition of transcriptional activity without affecting the cohesin loading (Cas9-ActD) results in a chromatin organization with increased heterochromatin domains sizes and LAD thickness, as shown in Figure 6b, top-right panel. On the other hand, the simulation of chromatin distribution in nucleus with depleted cohesin unloading, without disturbing the transcriptional activity, (WAPLΔ-DMSO) is shown in Figure 6b, bottom-left panel. Finally, the chromatin distribution predicted in a WAPLΔ nucleus with inhibited transcription (WAPLΔ-DMSO-treatment) is shown in Figure 6b, bottom-right panel. As reported in Section 3.3, ActD (mathematically, Γ_*tr*_ = 0 in Eq. 6) results in larger heterochromatin domains and thicker LADs, while WAPLΔ nuclei (increased cohesin loading; mathematically, Γ_*ul*_/Γ_*l*_ increases in Eq. 6) show the opposite effect with smaller heterochromatin domains and LADs. For a WAPLΔ nuclei in which transcription is inhibited (WAPLΔ – ActD; mathematically, Γ_*tr*_ = 0 and Γ_*ul*_/Γ_*l*_ increases in Eq. 6), the model predicts that inhibition of transcription returns the chromatin organization to the control (Cas9-ActD) levels. Transcription inhibition thus blocks the reduction in chromatin domain sizes induced due to WAPL deficiency due to lack of impetus for chromatin supercoiling.

#### In-vivo whole-genome organization validates the model predictions

To quantitatively validate the model predictions, we investigate the *in-vivo* chromatin organization under individual and tandem changes in transcription and cohesin unloading by re-analyzing previously reported super-resolution images shown as heatmap density plots in Figure 6c [14]. Visual inspection of this data agrees with the model predictions that transcriptional inhibition counteracts the chromatin blending observed in DMSO treated WAPLΔ nuclei, which was also previously reported [14]. We thus focused on extracting the radius of heterochromatin domains and LAD thickness to further validate the model results quantitatively (Figure 6d). Cas9 – ActD treated nuclei show an increased heterochromatin domain radius compared to control while WAPLΔ nuclei show a significant reduction in domain radius and LAD thickness (Figure 6d). However, WAPLΔ – ActD treated nuclei show no significant difference in comparison to Cas9 – ActD treated nuclei (Figure 6d), in quantitative agreement with the numerical predictions.

These results further confirm that the effect of transcription on global chromatin distribution occurs via the chromatin loop extrusion, especially at the interface of heterochromatin and euchromatin phases. Furthermore, these results also present a significant validation of the mathematical phase-field model of chromatin organization in the nucleus.

## 4 Discussion and Conclusions

While on one hand, spatial organization of chromatin into compacted heterochromatin domains and loosely packed euchromatin plays a significant role in governing gene expression, recent studies on a wide range of cell-types offer strong evidence to suggest that gene transcription via chromatin loop extrusion can in turn regulate the mesoscale organization of chromatin in the nucleus [2, 4, 14, 15, 35-39]. Molecular dynamics simulations have been used to predict the role of active RNAPII-mediated transcription in DNA supercoiling and chromatin loop extrusion at a nano-scale [15, 40-45]. However, quantitative predictions of sizes of heterochromatin domains which organize at a nucleus-wide meso-scale level are beyond the purview of such models. Furthermore, mathematical models for the genome-wide spatial organization [19, 46-49] lack the far from equilibrium dynamic considerations of active epigenetic regulation, chromatin extrusion and diffusion kinetics which we find are intricately involved in spatio-temporal size regulation of heterochromatin domains.

Here we present a non-equilibrium thermodynamic continuum model of the meso-scale chromatin organization in the nucleus to bridge the gap in the understanding of the mechanistic relation between transcriptional and epigenetic regulation and the size-scaling of the meso-scale heterochromatin domains. Our model incorporates the energetics of chromatin-chromatin interactions which is constructed as a double-well function allowing the phase-separation of chromatin into compartments of distinct compactions. Along the nuclear periphery, the effect of chromatin-anchoring proteins such as LAP2*β* is captured via energetic chromatin-lamina interactions leading to the formation of LADs. Concomitant with the energetics, the chromatin organization is temporally driven by diffusion kinetics of nucleoplasm and the epigenetic marks. While the diffusion of nucleoplasm determines the level of chromatin compaction, such that higher local nucleoplasm content results in lesser chromatin compaction, diffusion of epigenetic marks results in accumulation of acetylated and methylated nucleosomes driving their segregation (Figure S2, Section S1.5 of the Supplementary Information). Most importantly, we also account for the active reaction kinetics, which allow the interconversion of heterochromatin into euchromatin and vice-versa. The chromatin phase-interconversion can occur via the epigenetic regulation of chromatin in the nucleus via the acetylation or methylation of the histones. Finally, to capture the role of transcription mediated chromatin loop extrusion in the determination of the heterochromatin domain sizes, we incorporate a kinetic conversion of compacted chromatin into transcriptionally active euchromatin in presence of RNAPII.

Together the active transcriptional kinetics and epigenetic regulation determine the active interconversion of hetero- and euchromatin, thereby taking the chromatin organization in the nucleus to a dynamic steady-state configuration. Specifically, our theoretical analysis reveals that the active reaction kinetics alone – independently of energetic interactions – offer a complete control over the average extent of chromatin compaction in the nucleus, thereby breaking the detail balance of thermodynamic equilibrium. At the meso-scale, spanning individual heterochromatin domains, we theoretically observe that the distribution of epigenetic marks relevant to chromatin compaction exhibit a radial gradient which would drive an inward heterochromatin flux leading to ripening of the phase-separated domains. However, the presence of epigenetic and transcriptional regulation offers an opposition to the influx via – (a) acetylation of heterochromatin into euchromatin which is then pushed out via the diffusion of epigenetic marks, and (b) extrusion of loops of chromatin from the heterochromatin phase into euchromatin phase (refer Figure 2 e-g). The steady-state balance between the opposing fluxes leads to intrinsic emergence of heterochromatin domains of a characteristic size-scaling in our model. It is essential to note that without the active chromatin phase interconversion – at thermodynamic equilibrium – no inherent size-scale of heterochromatin domains would be observed.

Thus, our model predicts that transcriptional activity, synergistically with epigenetic regulation, controls the size and morphologies of the heterochromatin domains. The key predictions of the model as summarized in Figure 7c are:

**Figure 7.**
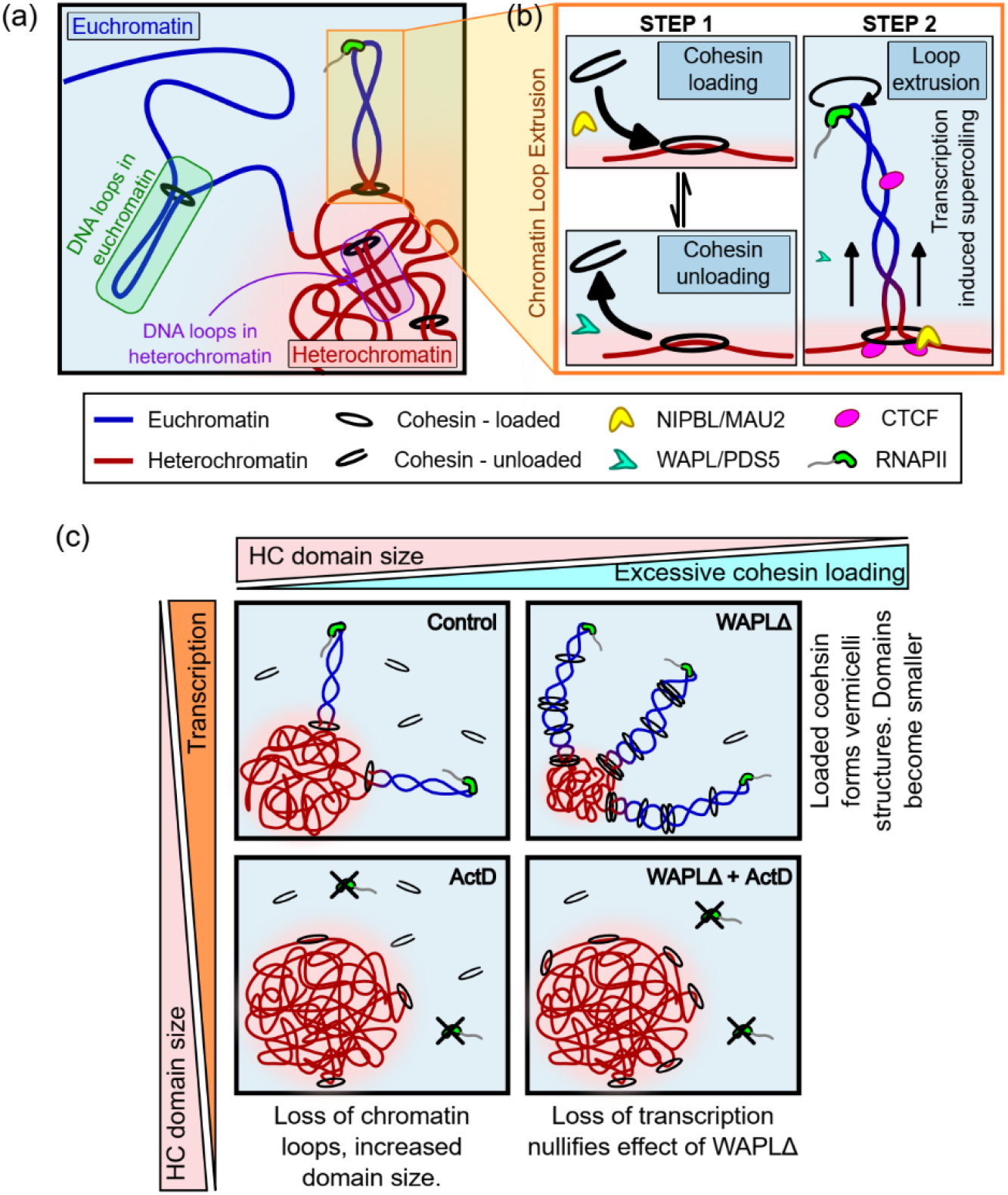
: A schematic summarizing the effect of chromatin loop extrusion on chromatin distribution in the nucleus. (a) A schematic of the roles of epigenetic and transcriptional regulation in chromatin compaction and (b) the steps in extrusion of a chromatin loop. (c) Schematics showing the alterations in chromatin compaction and loop extrusion upon inhibition of the proteins involved in the individual steps of loop extrusion.

1. Upon transcriptional inhibition, the characteristic sizes of heterochromatin domains increase due to loss of DNA loop extrusion (Figure 7c, left panels).
2. The increased size of heterochromatin domains upon transcription abrogation are also observed in the vicinity of the nuclear lamina.
3. Transcriptional inhibition leads to reduction of DNA in the euchromatic phase.
4. Conversely, upon increased loop extrusion due to excessive cohesin loading, the size of the heterochromatin domains reduces in the interior as well as periphery of the nucleus (Figure 7c, top panels).
5. Transcriptional inhibition in nuclei with excessive cohesin loaded, results in loss of loop extrusion resulting in increased domain sizes (Figure 7c, bottom right panel).

Being founded on fundamental non-equilibrium thermodynamic principles, the predictions made by our model are cell-type agnostic. Although the quantitative values of the parameters used to capture the energetic and kinetic phenomena, such as chromatin lamina interactions or rates of acetylation and methylation, might vary with cell types, our qualitative predictions are expected to hold across a wide range of cell lines. To validate this, complementary techniques such as Chrom-STEM (for high-resolution chromatin conformation imaging) and PWS (for high-throughput nano-scale sensitive live-cell imaging) are carried out for BJ fibroblast cells and epithelial cancer cell-lines – U2OS, HeLa, A549 and HCT116. We found that *in vivo* alterations in chromatin organization under transcriptional inhibition conditions are consistent with our model’s predictions across all studied cell lines. A quantitative analysis of previously reported [14] super-resolution STORM images of nuclei further gives a direct quantitative validation of the predicted effects of transcription abrogation on heterochromatin domain sizes. Of note, in all the reported cells, growth of condensed heterochromatin domains after ActD treatment is seen throughout the nucleus, including along the nuclear periphery where an increased LAD thickness is observed both *in-silico* and in cells. Lastly, our predictions on the changes in heterochromatin domain sizes upon over-extrusion of chromatin loops with and without transcription are quantitatively validated by domain size analysis of the previously reported [14] super-resolution STORM images of control and WAPLΔ nuclei after DMSO or ActD treatments.

Beyond transcription induced supercoiling, cohesin plays a role in formation, extrusion and maintenance of loops of chromatin at different physiological length scales. For instance, it has been reported that loss of cohesin results in a complete loss of loops identified as topologically associated domains (TADs) via chromatin contact mapping techniques [50]. Of note, here we are specifically interested in the meso-scale roles played by chromatin loops which extrude from the silenced heterochromatin phase into transcriptionally active euchromatin region (Figure 7a). These loops are specifically considered since the interconversion of heterochromatin to euchromatin will alter the sizes of the heterochromatin domains. Recent experimental evidence, as well as computational models, present strong evidence in favor of DNA loop extrusion mediated by the RNAPII driven transcription induced DNA supercoiling [4, 9, 14, 15, 36, 37, 51, 52]. Negatively supercoiled DNA regions are particularly rich in transcription start sites (TSS) with a strong correlation seen between transcription and supercoiling [32]. Indeed, super-resolution images show high presence of RNAPII at the heterochromatin-euchromatin phase boundaries where loops would extrude from heterochromatin into euchromatin phase [25].

The kinetics of chromatin loop extrusion from the compacted phase into transcriptionally active euchromatin is captured via a two-step process in our model (Figure 7b). First, cohesin is loaded onto a strand of DNA via the action of cohesin loaders such as NIPBL/MAU2. The transient loading is opposed by cohesin unloaders such as WAPL/PDS5. Cohesin loading is then followed by the transcriptional activity of RNAPII which induces negative supercoiling in the DNA thereby extruding loops of chromatin. These steps are incorporated via a first-order chemical reaction resulting in the non-conservative change of heterochromatin phase into euchromatin specifically localized to the chromatin phase-boundaries.

Intriguingly, previous observations [14] also show that in HeLa nuclei WAPL deficiency introduces abnormalities in the peripheral distribution of lamin A/C. Since lamin A/C plays an integral role in the chromatin-lamina interactions via chromatin anchoring proteins such as LAP2*β* and emerin, it can be conjectured that WAPL treatment may affect the LAD organization. In the current study we have ignored such effects focusing purely on the role of chromatin loop extrusion. Our model can be easily modified to address the LAD alterations by introducing WAPL deficiency dependent modulations in the chromatin lamina interaction parameter *V*_*L*_ in Eq 2. Experiment guided modifications in the model will further strengthen our predictions of LAD formation. Further, the transcriptional machinery involves a highly complicated multi-stage process comprising recruitment of multiple transcription factors, RNAPII and gene regulatory elements, we have assumed the cohesin loading and RNAPII mediated supercoiling to be the rate defining steps which thereby govern the timescale for chromatin loop extrusion. A more refined kinetic model of transcription and loop extrusion could possibly be incorporated to predict the spatio-temporal chromatin arrangement in the nucleus. However, even without these inclusions, we believe that our model lays a fundamental computational framework to better mechanistically understand the role of transcription, and in general chemo-mechanical cell-signaling, on the meso-scale chromatin organization.

## Acknowledgements

This work was supported by NIH Award U54CA261694 (V.B.S.); NCI Awards R01CA232256 (V.B.S.); NSF CEMB Grant CMMI-154857 (V.B.S.); NSF Grants MRSEC/DMR-1720530 and DMS-1953572 (V.B.S.); NIBIB Awards R01EB017753 and R01EB030876 (V.B.S.). This work made use of the BioCryo facility of Northwestern University’s NUANCE Center, which has received support from the SHyNE Resource (NSF ECCS-2025633), the IIN, and Northwestern’s MRSEC program (NSF DMR-1720139). We acknowledge the support from NIH grants U54CA268084 (V.B.), R01CA228272 (V.B.), R01CA225002 (V.B.), NSF grant EFMA-1830961 (V.B.), and philanthropic support from Rob and Kristin Goldman (V.B.). We acknowledge the support from the European Union’s Horizon 2020 Research and Innovation Programme (no. 964342 to M.P.C.); Ministerio de Ciencia e Innovación (grant no. 008506-PID2020-114080GB-I00 to M.P.C.; AGAUR grant from Secretaria d’Universitats i Recerca del Departament d’Empresa iConeixement de la Generalitat de Catalunya (grant no. 006712 BFU2017-86760-P (AEI/FEDER, UE) to M.P.C.).

## SUPPLEMENTARY INFORMATION

### S1 Extended Methods

#### S1.1 Mathematical description of chromatin distribution in nucleus

To investigate the organization of chromatin in the nucleus, we develop a mathematical model for the phase separation of heterochromatin and euchromatin considering chromatin-chromatin interactions, chromatin-lamina interactions, epigenetic regulation of chromatin via histone acetylation or methylation, and the role of transcriptional regulators. We consider three nuclear constituents – nucleoplasm and chromatin in either heterochromatin or euchromatin phases. At any point ***x*** in the nucleus, at a time *t*, the composition of the nucleus can be defined in terms of the local volume fraction of heterochromatin *ϕ*_*h*_(***x***, *t*), euchromatin *ϕ*_*e*_(***x***, *t*) and nucleoplasm *ϕ*_*n*_(***x***, *t*), such that *ϕ*_*e*_ + *ϕ*_*h*_ + *ϕ*_*n*_ = 1. Equivalently, the physical state of the nucleus at any point can be completely defined via two independent variables – (i) volume fraction of nucleoplasm *ϕ*_*n*_(***x***, *t*), and (ii) difference between the volume fractions of heterochromatin and euchromatin Φ_*d*_(***x***, *t*) = *ϕ*_*h*_(***x***, *t*) − Φ_*e*_(***x***, *t*). As discussed in Section 2.2 of Methods, *ϕ*_*d*_ can be considered an order parameter which when negative implies a euchromatin rich phase and when positive implies a more condensed heterochromatin rich phase. While the change of the variables is entirely equivalent mathematically, physiologically such a description permits a natural definition of the movement of the two mobile species in the nucleus – nucleoplasm or water, and the epigenetic marks of acetylation or methylation.

#### S1.2 Free energy landscape of the nucleus

In terms of the independent variables *ϕ*_*d*_(***x***, *t*) and *ϕ*_*n*_(***x***, *t*), the free energy density at any point ***x*** can be expressed as *W*(*ϕ*_*n*_, *ϕ*_*d*_, ∇*ϕ*_*n*_, ∇*ϕ*_*d*_) where we have also incorporated the energetic considerations associated with the phase interfaces via the spatial gradients of the volume fractions. Specific form of the free energy can be invoked by considering the various energetic contributions in the nucleus such as,

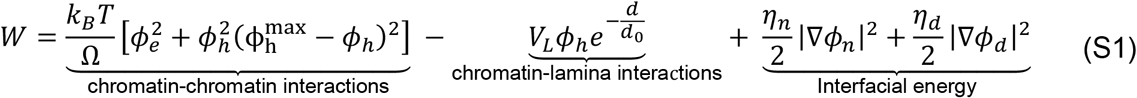

- The first term in Eq (S1) arises from the competition between entropy and enthalpy of mixing heterochromatin and euchromatin phases. It is equivalent to the Flory-Huggins free energy definition. This term gives rise to the double-well form of the free energy landscape, as shown in the contour plot in Figure 1b. The two wells, shown as black dots in the blue region, are the energy minima corresponding to the two stable phases of chromatin - a water-rich, loosely packed euchromatin phase (*ϕ*_*h*_ = 0) or a compacted water-devoid heterochromatin phase 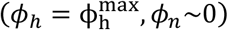. Here 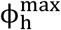 denotes the extent of compaction in the heterochromatin phase. Any initial chromatin configuration will spontaneously phase separate into heterochromatin and euchromatin domains (red arrows).
- The second term captures the interactions between the chromatin and the lamina via chromatin anchoring proteins (HDAC3, LAP2*β*, emerin, etc.) with parameter *V*_*L*_ denoting the strength of these anchoring interactions. These interactions are most robust at the nuclear periphery (distance from lamina *d* = 0) and vanish exponentially over a length scale *d*_0_. The negative sign permits an energetic preference for the heterochromatin phase along the nuclear periphery. We chose the heterochromatin phase specifically to interact with the lamina since the chromatin domains preferentially associating with the nuclear lamina are linked to transcriptional repression and an increased histone methylation [1-4].
- The last term in Eq (1) denotes the energy penalty associated with forming phase boundaries between the euchromatin and heterochromatin phases as they separate. The term *η*_*n,d*_ is the change in the energy due to formation of a unit width of the interface. Notably, as *η*_*n,d*_ increases, there is a greater penalty on formation of sharp interfaces, resulting in more smooth interfaces which are wider. Thus *η*_*n*_ and *η*_*d*_ directly control the width and the energy of the phase boundaries. In our simulations we choose *η*_*n*_ = *η*_*d*_ = *η*.

In Eq S1, *k*_*B*_ is the Boltzmann constant, T is the temperature and Ω is the volume of individual chromatin particles. Note that the coefficient *k*_*B*_*T*/Ω acts as an energy scaling factor. The total free energy of the nucleus can be written, after incorporating the work done in exchanging water between the nucleus and the cytoplasm, as,

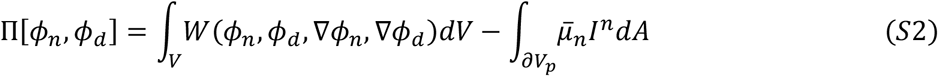

Here, *∂V*_*p*_ denotes the surface of the nucleus where the chemical potential of *I*^*n*^ is the volume of water entering per unit surface area into the nucleus, which determines the amount of water exchanged with the cytoplasm. The double-well energy landscape described by Eq (1) and Figure 1b, drive the time evolution (red arrows) of chromatin from an initial configuration (say corresponding to the red dot in Figure 1b) into the two energy minimal wells corresponding to the two chromatin phases. The gradients of the free energy in the *ϕ*_*n*_-*ϕ*_*d*_ variable space (shown by the contour plot in Figure 1b) provides the driving force for the time-evolution of chromatin organization towards the steady-state and is given by the chemical potentials obtained using variational principles as,

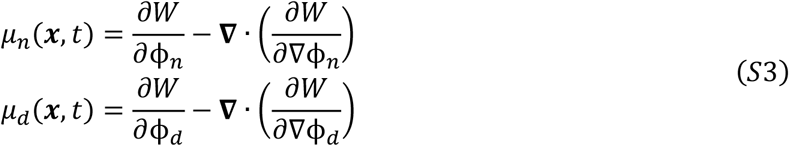

where, *μ*_*n*_ is the chemical potentials of nucleoplasm driving its kinetics as described in the next section. And *μ*_*d*_ is the chemical potential for the order parameter *ϕ*_*d*_ evolving the epigenetic marks in a conserved manner as discussed in the next sub-section.

#### S1.3 Diffusion and reaction kinetics

The dynamic evolution of the nucleus towards the steady-state is governed in a time-dependent fashion by a combination of diffusion and reaction kinetics (Figure 1b, bottom panel). The local conservation of reactively inert nucleoplasm content relates the time evolution of local nucleoplasm volume fraction with its nominal volumetric flux 𝒥^*n*^ as, 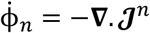. Fick’s first law gives the volumetric flux of nucleoplasm as 𝒥^*n*^ = −*M*_*n*_∇*μ*_*n*_ in terms of the gradient of the chemical potential of nucleoplasm where *M*_*n*_ is the mobility of water in the nucleus. Thus,

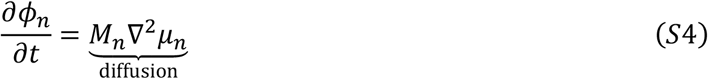

In a similar manner, the conservative change in the epigenetic marks between heterochromatin and euchromatin phases is driven by the gradient of the chemical potential *μ*_*d*_ such that,

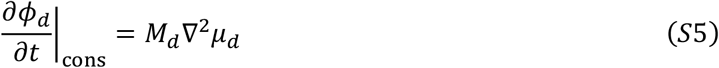

Here, *M*_*d*_ is the mobility of epigenetic marks in the nucleus. In addition, the non-conservative evolution of the epigenetic marks is facilitated by the epigenetic reactions.

**Fig S1:**
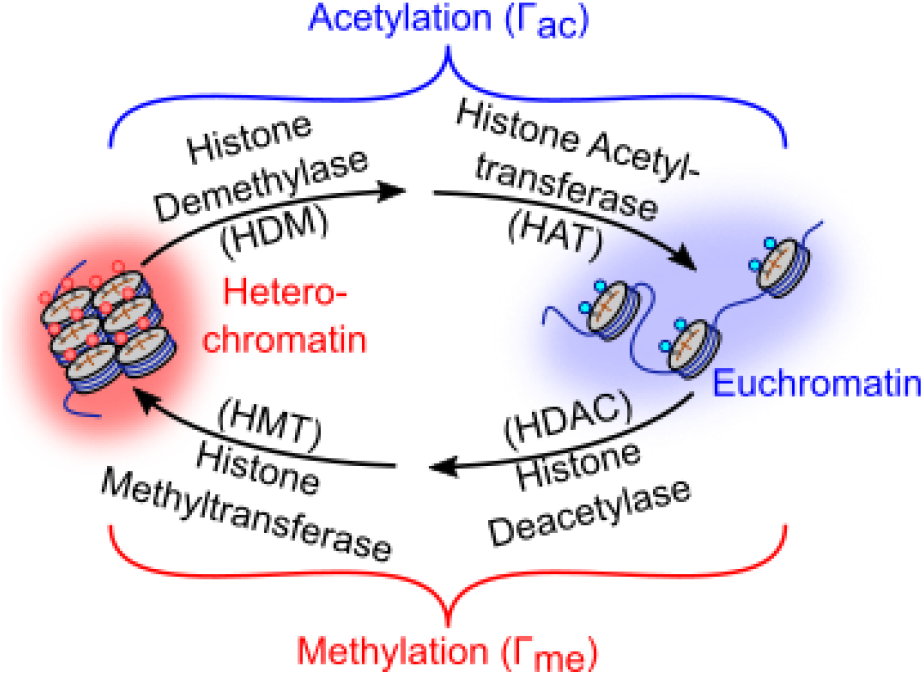
Epigenetic factors catalyze the reactions leading to interconversion of euchromatin and heterochromatin. The reactions are broadly methylation of euchromatin and acetylation of heterochromatin.

The epigenetic reaction kinetics controls the methylation or acetylation levels of the histone tail marks. Heterochromatin, which is rich in methylation marks on the histone tails, can be converted into euchromatin phase, where histones are marked by an increased acetylation level. This process encompasses the removal of methylation marks on the histone tails called demethylation via proteins classified as histone demethylase (HDM), followed by acetylation of the histone tails, via histone acetyltransferase (HAT) as shown in Figure S1. We classify both these processes together as the acetylation of the histone – occurring at a rate Γ_*ac*_ – which converts heterochromatin into euchromatin. Conversely euchromatin is converted into heterochromatin by first the deacetylation (via histone deacetylase, HDAC) followed by methylation (via histone methyltransferase, HMT) at a cumulative rate Γ_*me*_ as shown in Figure S1. Distinct from the conservative evolution in Eq (3), the interconversion of chromatin phases via epigenetic reactions changes the relative content of heterochromatin and euchromatin such that,

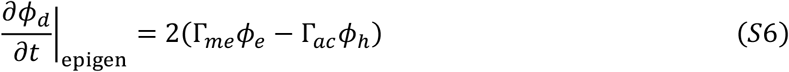

Lastly, active transcription requiring ATP-based energy expenditure, is known to supercoil chromatin fiber thereby resulting in active extrusion of DNA through cohesin rings. Since extruded chromatin loops are transcriptionally active, such extrusion within the euchromatin phase does not alter the gene expression. However, extrusion of transcriptionally silent heterochromatin into chromatin loops switches the transcriptional state of chromatin pulling out the silenced genes near the heterochromatin-euchromatin boundary into the euchromatin phase (Figure 1c). Chromatin extrusion at the phase boundaries happens in two steps (Figure 1c):

1. Cohesin rings entrap a portion of DNA fiber along the interface due to a balance between its loading (with reaction rate Γ_*l*_) via NIPBL/MAU2 and unloading (with reaction rate Γ_*ul*_) via WAPL/PDS5. The overall rate of cohesin loading can be written as Γ_*coh*_ = Γ_*l*_ − Γ_*ul*_.
2. This is followed by the active extrusion of supercoiled loops of chromatin through the cohesin rings via the transcription due to RNAPII, at a rate denoted by Γ_*tr*_.

Altogether, Γ_*a*_ = (Γ_*tr*_)^*α*^(Γ_*coh*_)^*β*^ denotes the overall the rate at which the chromatin extrusion converts heterochromatin into euchromatin via the two-step process (Figure 1c). Thus, we can write the transcriptionally-dependent conversion of the chromatin phases as,

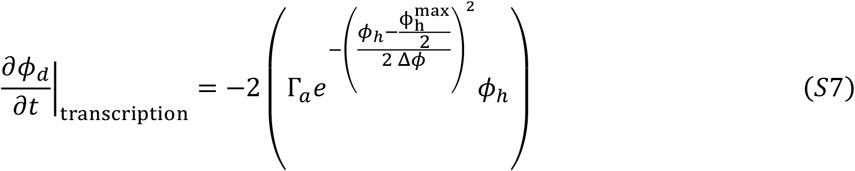

Where, the exponential factor ensures that the transcription-based chromatin extrusion is spatially restricted to a narrow region where the volume fraction of heterochromatin is *ψ*⁄2 − Δ*ϕ* ≤ *ϕ*_*h*_ ≤ *ψ*⁄2 + Δ*ϕ*, while peaking at the interface (*ϕ*_*h*_ = *ψ*/2). Combining Eq (4-6), the time-evolution of the order parameter *ϕ*_*d*_ can be written as,

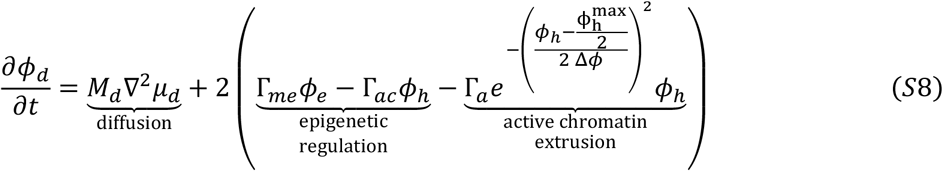

Eqs (S3), (S4) and (S8) together form the time-dependent mathematical set of governing equations describing the spatio-temporal evolution of chromatin organization in the nucleus.

#### S1.4 Rescaling the governing equations

The governing equations derived in the previous section reveal intrinsic length and time-scales which we next use to obtain the rescaled, non-dimensional set of governing equations.

The reaction-diffusion kinetics from Eq S8 results in a characteristic length scale determined by the reaction-diffusion kinetics i.e 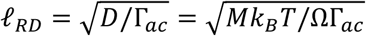. Interestingly, it can also be seen that another intrinsic length scale emerges from the competition between the interfacial and bulk mixing energies from Eq S1, i.e. the width of the interface 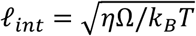. In our simulations, and from the theory discussed later in Section S4, we find that the reaction-diffusion length plays a significant role in determining the heterochromatin domain sizes and their spacing. Therefore, we choose to rescale all lengths with respect to *ℓ*_*RD*_, such that 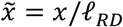. Further, the reaction rates offer an intrinsic time scale for the system of equations, such that all times are rescaled as 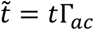. Lastly, *k*_*B*_*T*/Ω which is the coefficient of energy of chromatin phase interactions in Eq S1 provides the energy scaling such that all energy densities are written as 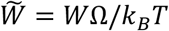.

Rescaling Eq S1,

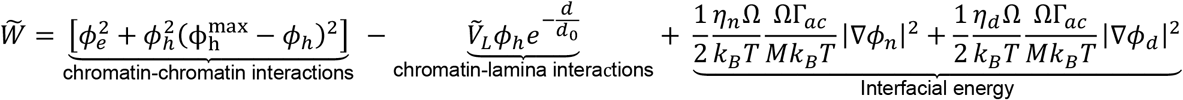

Here, 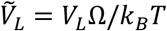 is the rescaled strength of chromatin-lamina anchoring interactions. Note that the coefficient of the interfacial energy terms can be rewritten in terms of the ratio of the length

Scales 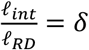. The parameter *δ* is a rescaled measure of the width of the interface. Thus,

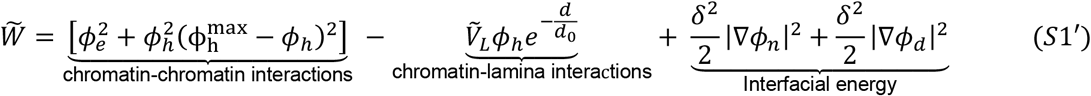

The rescaled chemical potentials from Eq S3 are,

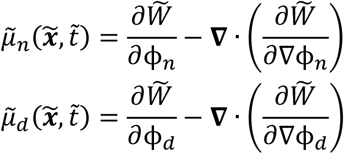

Expanding these,

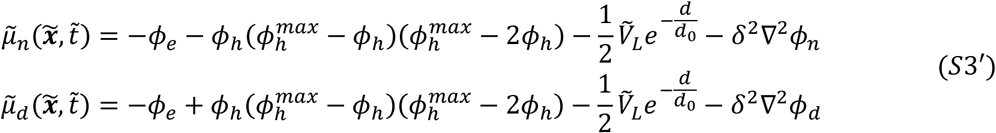

Lastly, we rescale the kinetics equations such that,

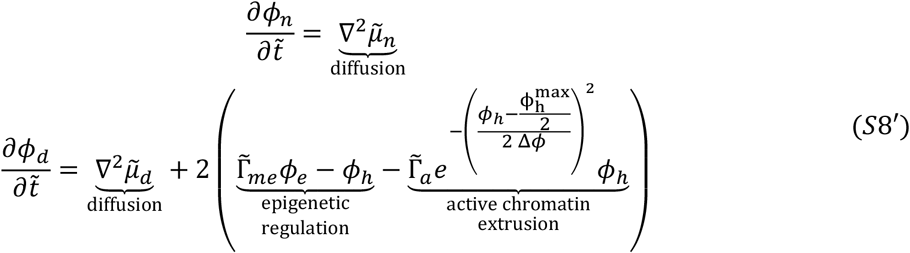

Note that in the second part of Eq S8’, all reaction rates have also been rescaled with respect to the time scale such that 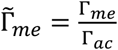 and 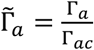. To solve the rescaled set of governing equations, Eq S3’ and Eq S8’ are solved together subjected to boundary conditions of no flux of nucleoplasm or epigenetic marks across all boundaries.

#### S1.5 Polymer analogy of the roles played by reaction and diffusion kinetics

As discussed in the previous section, we have incorporated the kinetics of both diffusive and reactive nature – the former being conservative, i.e. it does not change the net amount of heterochromatin and euchromatin in the nucleus, while the latter non-conservative since it allows interconversion of the two phases (Figure S1). Here we explain in detail the effects of non-conservative and conservative kinetics.

The diffusion kinetics given by Eq (S4) and (S5) are conservative in nature as these equations keep the amount of nucleoplasm (*ϕ*_*n*_) and the order parameter denoting the epigenetic marks (*ϕ*_*d*_) constant within the nucleus, as long as there is no flux occurring across the nuclear lamina. The conservation of nucleoplasm via Eq (S4) dictates that the total number of water molecules in the nucleus stays the same although they may relocate within the nucleus, as shown in Figure S2a. Similarly, conservation of epigenetic factors via Eq (S5) dictates that the total amount of DNA in heterochromatic or euchromatic phase in the nucleus does not change, although methylated or acetylated histones may move along the chromatin polymer (Figure S2b).

**Fig S2:**
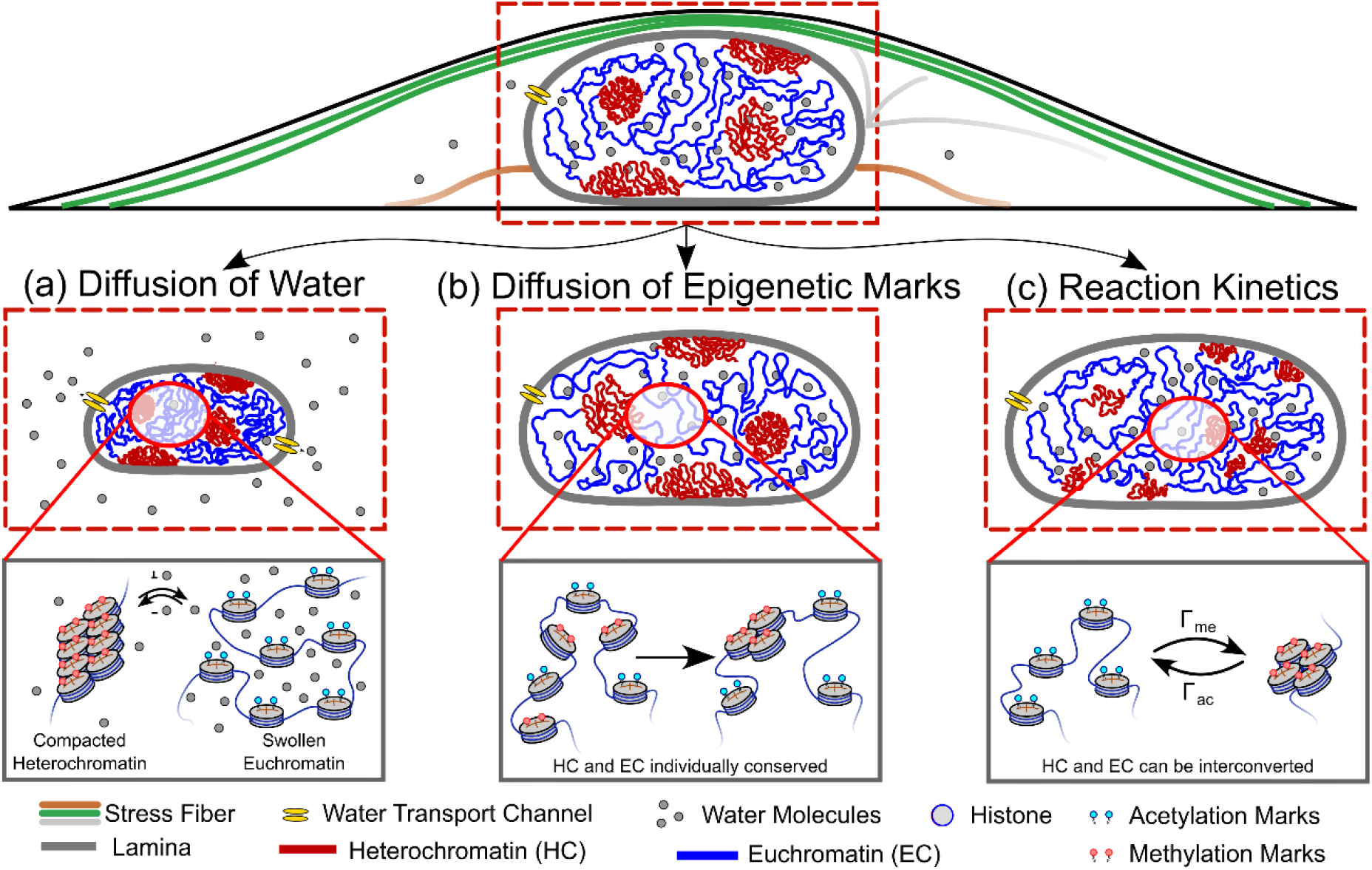
A schematic depicting the individual roles of the diffusion and reactions kinetics incorporated into the heterochromatin organization model. Column 1 denotes the conservative diffusion of water which can redistribute molecules of water within the nucleus without changing the total amount of water, and consequentially the total amount of DNA, in the nucleus. Column 2 denotes the conservative diffusion kinetics of epigenetic marks of methylation and acetylation. This may allow a relocalization of methylation or acetylation histones without changing the overall amounts of heterochromatin and euchromatin. The last column denotes the non-conservative reaction kinetics of histone acetylation and methylation, which allows an interconversion of chromatin phases. This changes the individual amounts of heterochromatin and euchromatin in the nucleus without changing the total amount of DNA. The reaction rates determine the ratio of heterochromatin to euchromatin at steady state.

The epigenetic reaction kinetics, on the other hand, as given by Eq (12) is non-conservative in nature as it allows an interconversion of euchromatin to heterochromatin via a reaction rate (or analogously a probability) Γ_*me*_ and heterochromatin to euchromatin via a reaction rate Γ_*ac*_. While this does not affect the total amount of water or DNA in the nucleus, it changes the individual amounts of heterochromatin and euchromatin. In other words, there may be a movement of acetylation marker (or methylation marker) from one histone to another. The exchange of heterochromatic and euchromatin phases driven by the non-conservative epigenetic reaction kinetics is shown in Figure S2c.

In addition to the above-mentioned kinetics, in the main manuscription we discuss in detail the role played by the reaction kinetics governing the transcription mediated active extrusion of chromatin loops Γ_*a*_ (Figure 1c, Eq 13). In short, the active extrusion kinetics is also a non-conservative kinetics that allows the conversion of heterochromatin phase into euchromatin phase. However, as dictated by the spatial dependence of Γ_*a*_ given in Eq (13), the chromatin extrusion is restricted to only the phase boundaries (discussed in Section 2.1, Section 4).

#### S1.6 Chromatin clustering of STORM images

MATLAB was used for the analysis of STORM images. For chromatin density qualification, Voronoi tessellation-based segmentation was implemented to construct Voronoi polygon of each localization [5]. The corresponding Voronoi polygon of each locus represents a specific region where any points within this region are closer to this locus, such that the size of Voronoi polygon is inversely proportional to the local Voronoi density. Voronoi polygons located at the edge were omitted due to nearly infinite Voronoi area. The Voronoi density map was further constructed through calculating the reciprocal of Voronoi area map. To differentiate between hetero- and eu-chromatic regions, a density-based threshold was applied to filter out low density euchromatic region. The threshold density value was chosen such that 60∼100 percentile of the Voronoi density distribution were classified as heterochromatin in the control group. The same density threshold was applied to all other treatments in comparison with the control. The remaining heterochromatin points cloud was then clustered using Density-based spatial clustering of applications with noise (DBSCAN) algorithm [6]. The heterochromatic point clouds were clustered into separated subdomains, such that each subdomain includes all the neighboring and connected Voronoi polygons which are above threshold. The hyperparameters of DBSCAN were set such that the minimal scan points are greater or equal to the dimension of dataset plus 1, namely 3 in our case [6].

#### S1.7 Quantitative analysis of chromatin distribution in STORM images

Chromatin clusters obtained after the DBSCAN subdomain classification were categorized as LADs and non-LADs depending on proximity to the boundary of the nucleus image. The nucleus shape was detected using boundary detecting algorithm. A characteristic radius of nucleus (R) was then calculated. The minimal distance between each heterochromatin subdomain and nucleus boundary was calculated, such that any subdomain having a distance smaller than 0.15*R* is classified as LADs domain [7].

To quantify the size of non-LAD heterochromatin subdomains, the area of each non-LAD domain was calculated through detecting the boundary of point clouds which gives a polygon enveloping it. An approximating domain radius was calculated by assuming the subdomain to be a circular shape 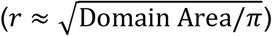.

The local LADs thickness was measured by sampling the LAD boundary along the nucleus periphery.

#### S1.8 ChromSTEM sample preparation, imaging, and reconstruction for BJ Fibroblasts

BJ cell lines (ATCC Manassas, VA) were cultured in Minimum Essential Media (ThermoFisher Scientific, Waltham, MA,#11095080) at physiological conditions (5% CO 2 and 37 °C). Cells were seeded on 35-mm glass-bottom Petri dishes (MatTek Corp.) until approximately 40-50% confluent and were given at least 24 hours to adhere to the dish before fixation. For ChromSTEM sample preparation, the previously published protocol was adapted [8]. Before fixation, cells were thoroughly rinsed three times in Hank’s balanced salt solution without calcium and magnesium (EMS). Cells were fixed using 2.5% EM grade glutaraldehyde, 2% paraformaldehyde, 2 mM CaCl_2_ in 0.1 M sodium cacodylate buffer, pH = 7.4 at room temperature for 5 minutes and then replaced with fresh fixative and fixed on ice for an hour. The cells were then washed with 0.1 M sodium cacodylate buffer 5 times on the ice. The cells were then incubated in a blocking buffer (10 mM glycine, 10 mM potassium cyanide in 0.1 M sodium cacodylate buffer, pH = 7.4) for 15 minutes, followed by staining with 10 μM DRAQ5™ (Thermo Fisher) and 0.1% saponin solution in 0.1 M sodium cacodylate buffer, pH = 7.4 for 10 minutes. After washing with the blocking buffer twice, the cells were incubated in the blocking buffer on ice before photo-bleaching. During photobleaching on a cold stage using continuous epi-fluorescence illumination (150 W Xenon Lamp) with Cy5 red tilter with a 100x objective for 7 minutes, the cells were incubated in 2.5 mM of 3–5′-diaminobenzidine (DAB) solution (Sigma Aldrich) in 0.1 M sodium cacodylate buffer, pH = 7.4. The cells were washed with 0.1 M sodium cacodylate buffer five times and then stained with reduced osmium solution (EMS) containing 2% osmium tetroxide, 1.5% potassium ferrocyanide, 2 mM CaCl_2_ in 0.15 M sodium cacodylate buffer, pH = 7.4 for 30 minutes on ice. Then the cells were washed with double distilled water five times on ice. Serial ethanol dehydration (30% ethanol, 50%, 70%, 85%, 95%, 100%x3) was followed by Durcupas resin (EMS) infiltration. An infiltration mixture containing equal proportions of 100% ethanol and Durcupan ™ resin mixture 1 (10 mL Durcupan ™ ACM single component A, M, epoxy resin, 10 mL Durcupan ™ ACM single component B, hardener 964, and 0.15 mL Durcupan ™ ACM single component D) was used to infiltrate cells for 30 minutes at room temperature. Next, an infiltration mixture containing 5 mL 100% ethanol and 10 mL Durcupan ™ resin mixture 1 was used to infiltrate the cells for 2 hours at room temperature. Durcupan ™ resin mixture 2 (0.2 mL Durcupan ™ ACM, single component C, accelerator 960 to mixture 1 (10 mL of component A, 10 mL of component B, and 0.15 mL of component D) was used to infiltrate the cells at 50°C in the dry oven for 1 hour. The photobleached cells were embedded flat with Durcupan ™ resin mixture 2 in beem capsules and further cured at 60 °C in the dry oven for 48 hours. An ultramicrotome (UC7, Leica) was used to section 100 nm thick slices that were deposited onto a copper slot grid with carbon/Formvar film. Then, 10 nm colloidal gold fiducial markers were carefully deposited on both sides of the sample. A 200 kV cFEG STEM (HD2300, HITACHI) with HAADF mode was used and while keeping the field of view constant, the sample was tilted from − 60° to 60° with 2° increments on two roughly perpendicular axes. The fiducial markers were used to align the tilt series in IMOD [9] and reconstructed using Tomopy [10] with a penalized maximum likelihood for 40 iterations independently.

#### S1.9 Domain Center Mapping and Statistical Analysis

The centers for individual chromatin domains were estimated from local maxima obtained from ChromSTEM projection with enhanced contrast in FIJI as previously described [11]. Domains occupying less than 50% volume in the z-plane were excluded from the analysis as they could be incomplete parts of other neighboring domains. Mass scaling analysis and radial density analysis were then performed originating from the identified individual domain centers. Average mass scaling originating from the individual domain centers was estimated using the area (mass) weighted by the grayscale ChromSTEM intensity within concentric circles with increasing distances from the domain centers. Similarly, radial chromatin density was estimated as the grayscale ChromSTEM intensity within concentric circles with increasing distances from the domain centers (Figure S3). We have shown that the radial mass density originating from the domain center decreases with increasing distance and approaching the domain boundary and then increases as the boundary of neighboring domains starts interacting. Both the mass scaling behavior and the radial chromatin density profile for each domain were then utilized to obtain the boundary or the approximate radius of the domain. The mass scaling approximately follows power-law scaling up to a given length scale and can be represented by a given slope or scaling exponent, *D* based on the linear regression of the mass scaling curve in the log-log scale. Beyond the domain regime, the slope gradually increases to reach the supra-domain regime. The radius of chromatin packing domains was estimated as the smallest length scale where the mass scaling curve deviates from the initial power law calculated from small length scales by 5% and the radial chromatin density starts to increase after a gradual decrease. The distributions for packing domain radius and density are shown as mean ± S.D. using violin super plots [12].

**Fig S3:**
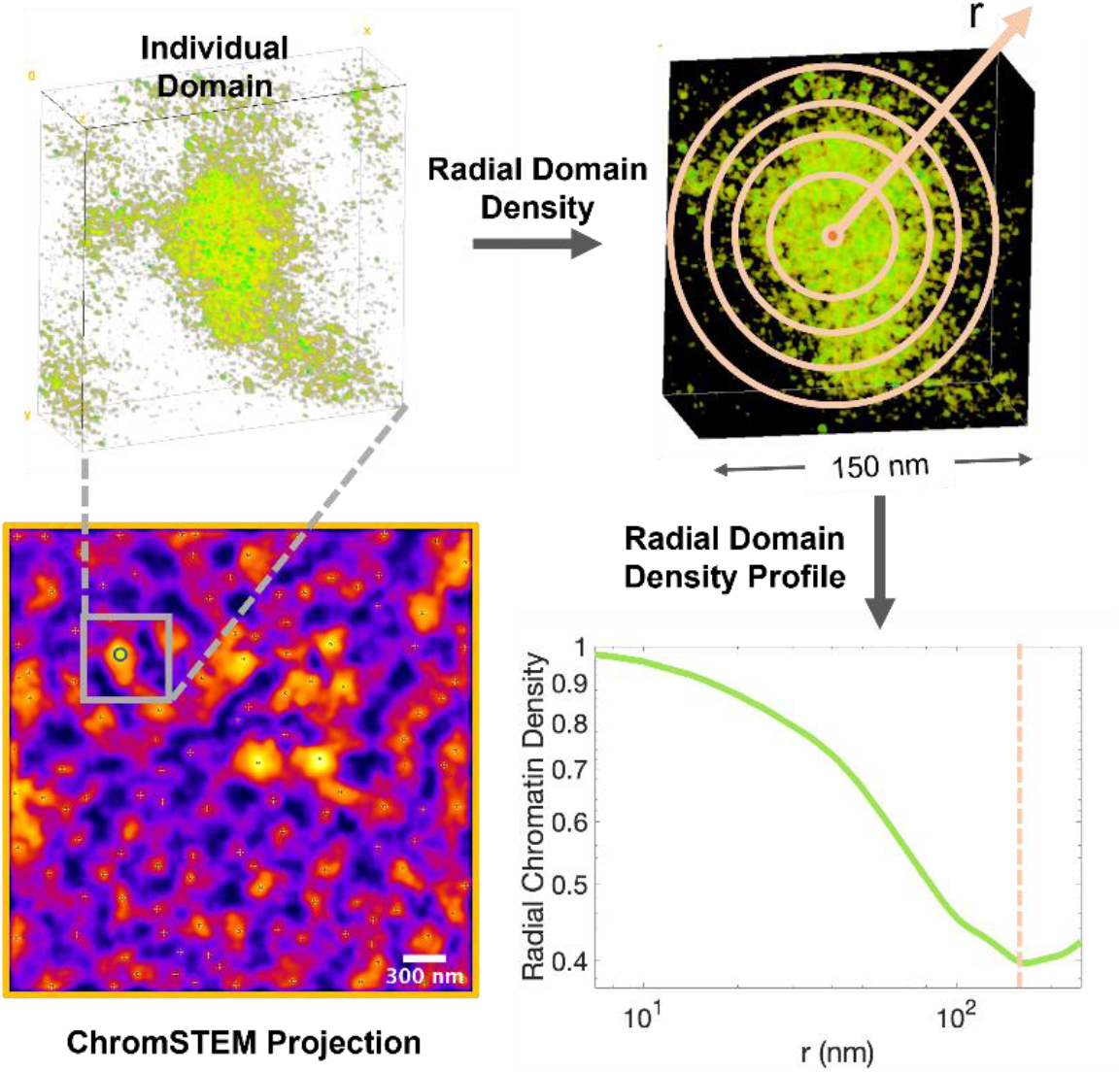
Radial density analysis to establish the radial profile of chromatin packing density estimated as grayscale ChromSTEM intensity within concentric circles with increasing distances from the domain center.

#### S1.10 ActD Treatment for PWS and ChromSTEM imaging

Prior to imaging, cells were cultured in 35-mm glass-bottom Petri dishes until approximately 70% confluent. All cells were given at least 24 hours to re-adhere before treatment (for treated cells) and imaging. HCT116, A549, HeLa, and U2OS cells were treated with Actinomycin D (Gibco, Cat: 11805017) for 1 hour at a final concentration of 5 μg/ mL.

#### S1.11 PWS Image Acquisition and Approximation of domain size scales from PWS image analysis

For live-cell measurements, cells were imaged and maintained under physiological conditions (5% CO2 and 37°C) using a stage-top incubator (In Vivo Scientific, Salem, SC; Stage Top Systems).

The PWS optical instrument was built on a commercial inverted microscope (Leica, Buffalo Grove, IL, DMIRB) supplemented with a Hamamatsu Image-EM CCD camera C9100-13. This camera was coupled to an LCTF (CRi Woburn, MA) for hyperspectral imaging. Spectrally resolved images of live cells were collected between 500 and 700 nm with a 2-nm step size. Broadband illumination was provided by an Xcite-120 light-emitting diode lamp (Excelitas, Waltham, MA). PWS is a high-throughput, label-free approach that measures the spectral standard deviation (Σ) of internal optical scattering originating from nuclear chromatin. The variations in the refractive index distribution Σ, are characterized by a mass density autocorrelation function (ACF) to calculate chromatin packing, scaling *D*.

Based on ChromSTEM [11], we have previously reported that chromatin packs into domains, wherein each domain exhibits a polymeric fractal-like behavior and can be described by an average packing scaling exponent. This implies that within the fractal regime, the genomic size of chromatin scales with its physical size following a power law relationship. Therefore, we estimated the upper bound of the power law regime as a measure of domain size.

Thus, a power-law ACF which incorporates a lower and upper length scale limit of the power law regime was utilized for the subsequent approximations,

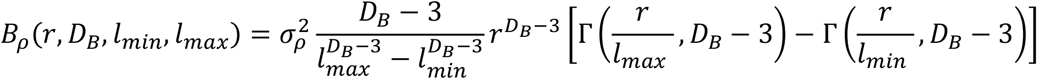

where Γ(*x,a*) is the upper incomplete gamma function, and *l*_min_ and *l*_max_ characterize fractality’s lower and upper length scales, respectively; *B*_*ρ*_(*r*=0) is *σρ*^2^, the variance of chromatin mass density; *D*_*B*_ describes the shape of *Bρ* and is related to *D*, and *r* is the spatial separation Utilizing this previously described methodology [13], we evaluated *l*_max_, the upper length scale of chromatin mass density scaling to estimate the relative size of domains upon ActD treatment.

### S2 Heterochromatin domain morphology dependence on Epigenetic Rates

**Fig S4:**
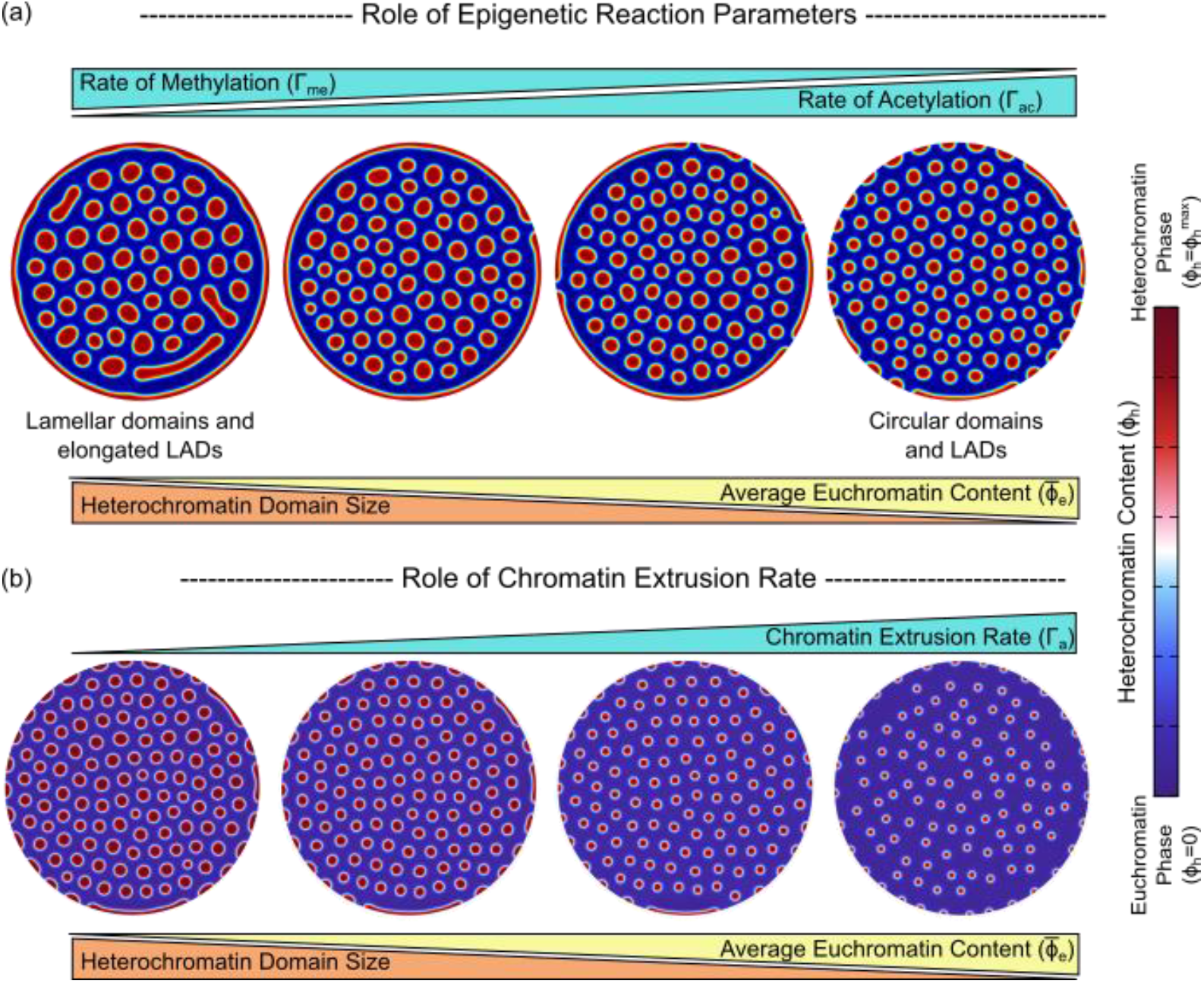
A study of the effect of reaction parameters (a) for epigenetic regulation and (b) rate of chromatin extrusion governed by transcription.

Our model predicts that the heterochromatin domains obtained at the steady state display a characteristic size. The length scale of the stable domains is regulated in tandem by the epigenetic reactions – acetylation as well as methylation – and the transcriptionally active extrusion of chromatin loops. As the levels of histone acetylation is increased (or conversely methylation is decreased), as shown in Figure S4a, we see that the sizes of the heterochromatin domains decrease. This trend is displayed not just by the interior heterochromatin domains, but also by the LADs near the nuclear periphery.

Also note that the morphology of the heterochromatin domains is regulated by the balance of acetylation and methylation reaction kinetics. Under the conditions where acetylation predominates over methylation, the domains formed are more circular. On the other hand, when methylation is increased while acetylation is reduced, the domains become larger and predominantly lamellar.

In addition to methylation and acetylation, transcriptional activity also regulates the sizes of the heterochromatin domains (Figure S4b). This is discussed expansively in Section 2 and 3 of the main manuscript (also see Figure 2c).

### S3 Theoretical analysis of average chromatin phase contents determined by reactions

Here we show that the total amount of chromatin which falls into the individual phases, i.e. either euchromatin or heterochromatin, can be shown to be determined solely by the epigenetic and chromatin extrusion reaction kinetics. The spatio-temporal evolution of order parameter *ϕ*_*d*_ due to the presence of diffusion of epigenetic marks and reaction driven interconversion of eu- and heterochromatin phases, as described by Eq 2 (or equivalently, Eq S8) as,

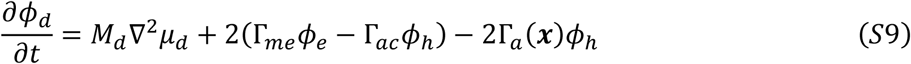

Once steady state is reached, the evolution halts and 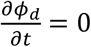. At this stage, the relation between the average euchromatin content 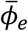 and the average heterochromatin 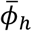 content in the nucleus can be obtained by averaging Eq 9 (i.e., integrating over the entire nucleus or region of interest and divide by the area of the nucleus) as,

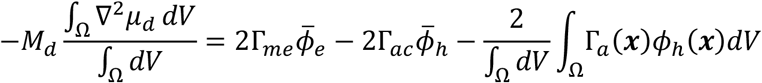

Note that the term ∇^2^*μ*_*d*_ on the left-hand side is non-zero near the interface and after averaging over the entire volume can be approximated to zero. Thus,

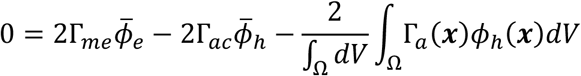

The last term on the right-hand side can be resolved via integration by parts as,

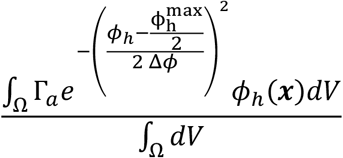

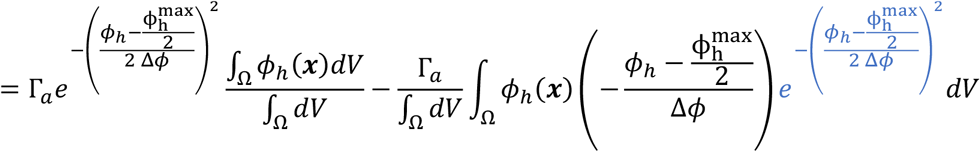

The last term (in blue) when integrated over the domain measures the length of the interfaces between the heterochromatin and euchromatin domains. The term in red is non-zero only along the interface. For a narrow interface width Δ*ϕ* → 0, this can be approximated as multiplying the integrand by a factor depending on the length of the interface giving a parameter *ℓ*_*int*_. Thus,

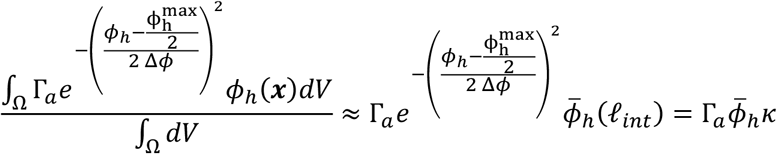

Thus, the equation can be written as,

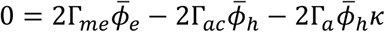

*k* being a non-trivial function dependent on 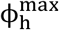, volume fraction change across the interface Δ*ϕ*, and the length of the interface between the two chromatin phases. By definition of the volume fractions, *ϕ*_*e*_ = 1 − *ϕ*_*n*_ − *ϕ*_*h*_. Thus,

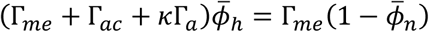

Or,

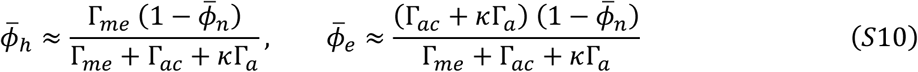

In the absence of transcription (Γ_*a*_ = 0), we can instead write the average heterochromatin and euchromatin contents as,

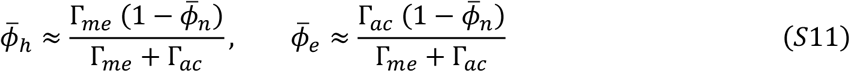

### S4 Domain size determination in presence of transcription – theoretical analysis

**Fig S5:**
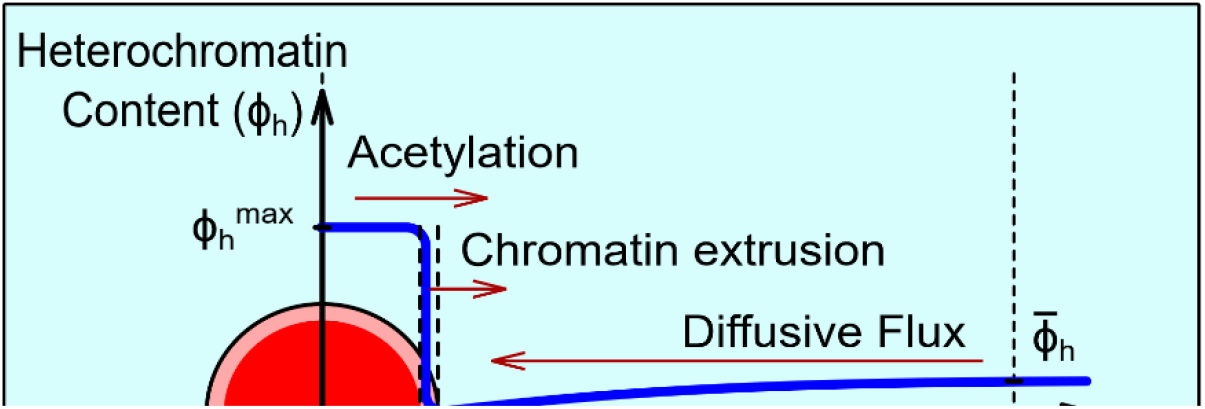
The competition of diffusion driven influx of heterochromatin with the epigenetic reaction and transcription mediated extrusion driven outflux of heterochromatin from the heterochromatin domain determines its steady state size. The figure also shows the radial distribution of heterochromatin volume fraction *ϕ*_*h*_ in and around the domain.

The steps involved in theoretical derivation of heterochromatin domain size have been enumerated in the main text (Section 2.1). Here we show the complete derivation of domain size determination in the interior of the nucleus, away from the periphery. To analyze the steady state size of the heterochromatin domain, we first examine the volume fraction fields within and around the droplet. As discussed in Section S3, the acetylation, methylation and chromatin extrusion together determine the mean heterochromatin (and euchromatin) volume fraction in the nucleus, given by Eq S10. However, a homogenous mean chromatin composition 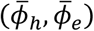 lies in neither of the energy wells as shown in Figure 2b (light blue circle) and is thus energetically unfavorable. As the system marches towards a steady state, its free energy must reduce, requiring the phase separation to initiate via nucleation of heterochromatin droplets (Figure S5).

Under a dilute limit, i.e., when there is a lot more euchromatin than heterochromatin, we can assume that the droplet size is much smaller than the length scale of the interdomain spacing such that neighboring droplets are far enough to not interact with each other. Under such assumption the heterochromatin distribution would be spherically symmetric. Using a polar coordinate system with origin at the droplet center, we can determine radial distribution of the heterochromatin volume fraction *ϕ*_*h*_(*R*) as,

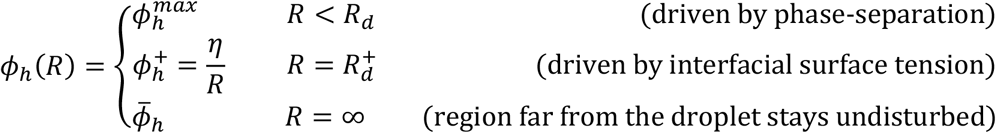

where *R*_*d*_ is the rescaled radius of the droplet at the current instance, while *η* measures the interfacial energy. Figure S5 shows the distribution of heterochromatin volume fraction *ϕ*_*h*_(*R*) around a single spherical domain of heterochromatin (red) of radius *R*_*d*_ as it grows surrounded by euchromatin phase (in blue). The slope of the heterochromatin profile outside the droplet will drive an inward flux of heterochromatin into the droplet. The volume fraction field outside the droplet at steady state must follow the equation,

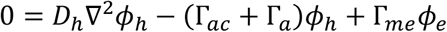

with boundary conditions 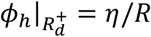 and 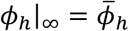, and thus must have the form,

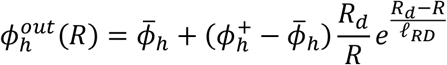

where *ℓ*_*RD*_ is the characteristic reaction-diffusion length scale given under a dilute limit as 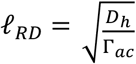. Note that the transcription driven active extrusion occurs only at the periphery and thus does not play a role in the reaction-diffusion length scale. Thus,

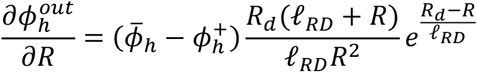

Thus,

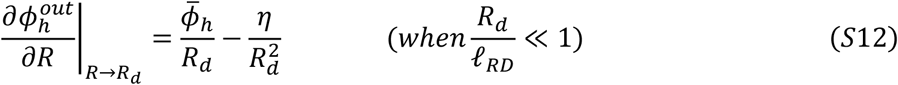

Thus, at the periphery of the droplet, the diffusive influx of heterochromatin into the droplet due to the reaction-diffusion phenomena outside is,

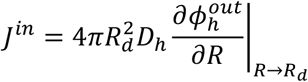

The inward diffusion is opposed by the outward flux of heterochromatin into euchromatin phase which occurs due to both acetylation of histones inside as well as DNA loop extrusion along the domain periphery. Thus, the rate of change of the volume of the droplet *V*_*d*_ can be written as,

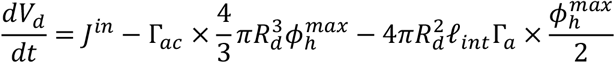

where *ℓ*_*int*_ is the width of the interface between the chromatin phases. Simplifying this we obtain,

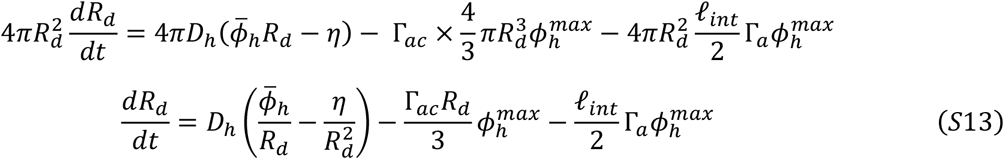

Equation S13 is reproduced in the main text as Eq 4 for a small value of interfacial energy. Using Eq S13 to plot the rate of change of heterochromatin domain size with respect to the instantaneous domain radius we obtain the plot shown in Figure S6.

**Fig S6:**
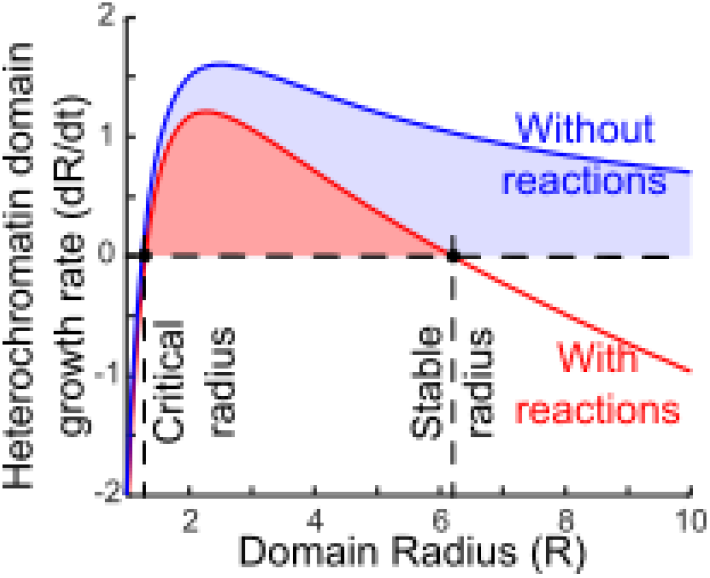
The growth rate of the heterochromatin domain varying with its instantaneous radius.

Above a critical radius, all heterochromatin domains growth in size (*dR*_*d*_/*dt* > 0). In the absence of reactions (Γ_*ac*_ = Γ_*a*_ = 0, blue curve), we note that the rate of increase in the heterochromatin domain radius is always positive indicating that the domain will keep growing as long as its radius is larger than the critical radius. However, in the presence of the reactions, we note that the domains will grow until their growth rate reaches a zero value. This gives the stable size of heterochromatin domains i.e. domains which neither grow nor shrink. The domains larger than the stable radius will shrink back to the stable radius.

The stable radius can be obtained by setting *dR*_*d*_/*dt* = 0 in Eq S13 such that,

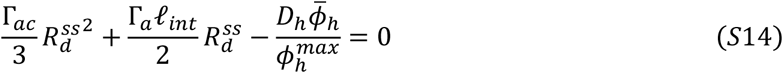

Thus,

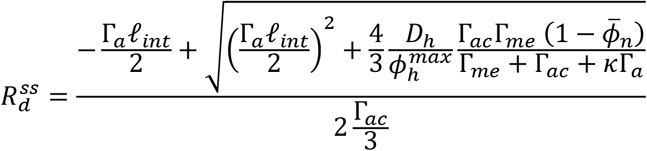

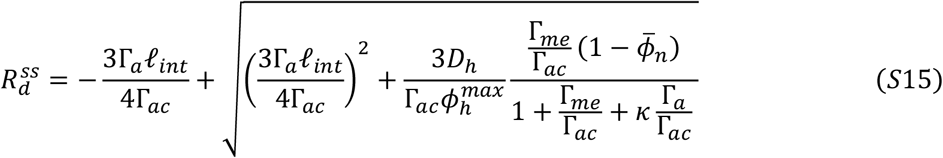

Note that in the absence of transcription, the steady state domain size can be obtained by substituting Γ_*a*_ = 0 as,

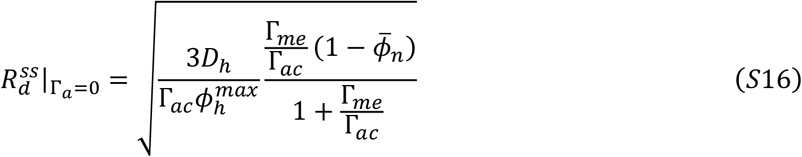

Note that the interior heterochromatin radii in the nuclei treated with ActD must approximately follow Eq S16. Further, we have assumed a dilute limit for the analytical derivations, which implies that the neighboring heterochromatin domains do not interact with each other. Such assumption therefore requires that the domains be separated by a distance which scales with the reaction-diffusion length scale such that the spacing between the domains can be written as,

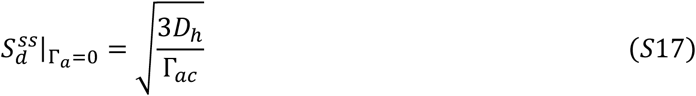

Thus, a quantitative image analysis of super-resolution images of heterochromatin foci in in-vitro nucleus can be used to quantitatively estimate the parameter Γ_*me*_ using Eq S16. As discussed in Section S6, we will use these relations to motivate the parameter choice for our numerical simulations.

We can rescale Eq S15-17 to obtain a non-dimensional dependence of heterochromatin domain size on the epigenetic and transcriptional kinetics. All lengths are rescaled with respect to *ℓ*_*RD*_, while all times with respect to 1/Γ_*ac*_. Thus, Eq S15 becomes,

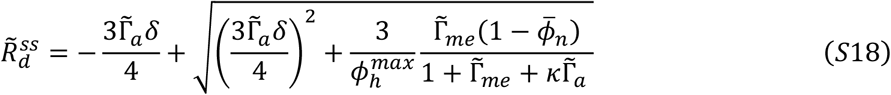

While Eq S16 and S17 become,

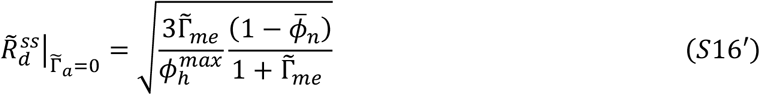

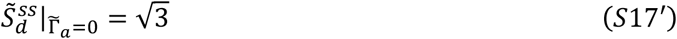

### S5 A characteristic size of heterochromatin domains is not obtained without reactions

As discussed in Section 2.1, the steady state organization of chromatin the nucleus (Figure 2a) comprises of many disconnected domains of heterochromatin phase 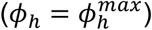 with a characteristic size. We also determined that the size of these domains is determined by the reaction kinetics – the rates of acetylation Γ_*ac*_, methylation Γ_*me*_ and active chromatin extrusion Γ_*a*_ (Eq 5).

To investigate the role of reactions numerically, we allow the phase-separation to occur from the same initial state as in Figure 2a, but without any reaction kinetics. The initial, intermediated and steady state chromatin organization thus obtained is shown in Figure S7a. While intermediate steps show the nucleation of multiple domains, as the organization evolves these domains merge. At the steady state, a single domain of heterochromatin remains. The growth rate of the domains given by Eq (4) is plotted in Figure S7b. In the absence of reactions, *dR*/*dt* never goes to zero, except at the critical radius. The critical radius only ensures that the domains above this size grow, while the rest shrink. Since no stable radius is predicted, the growing domains continue to grow 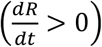, until all heterochromatin merge into a single domain.

**Fig S7:**
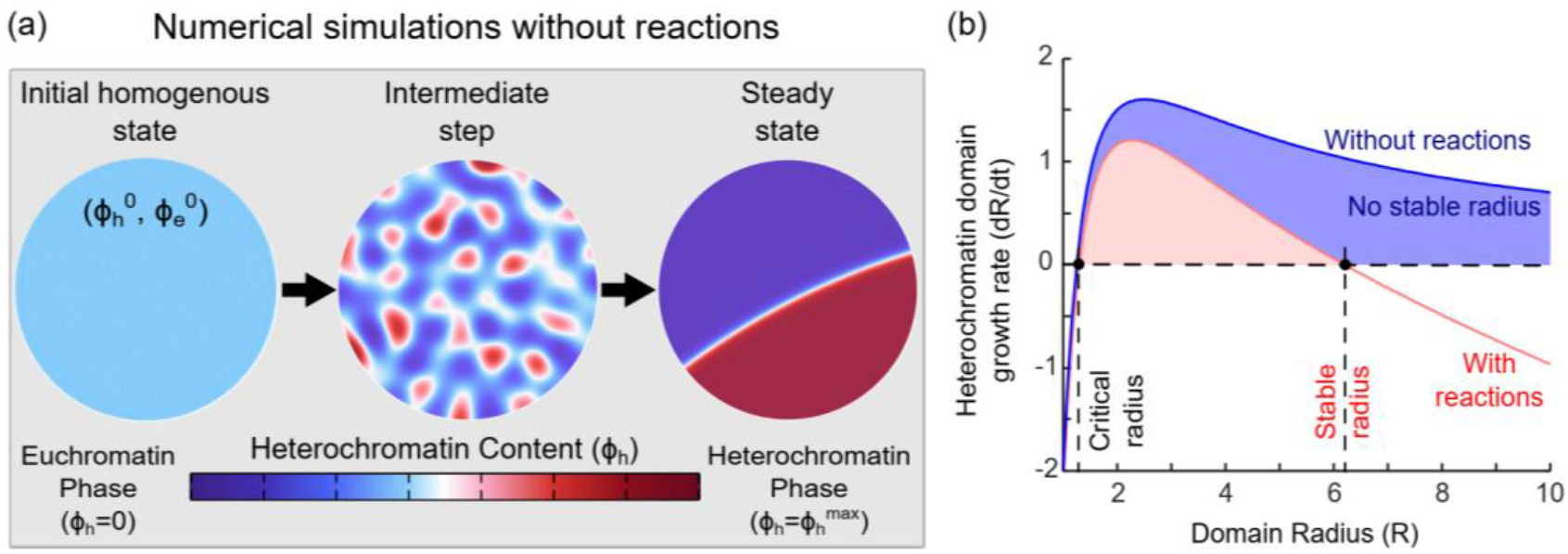
Steps in the numerical simulation show the evolution of chromatin organization in absence of the epigenetic and transcriptionally mediated reaction kinetics. While nucleation of multiple domains does occur, as they evolve all the nucleated domains merge into a single heterochromatin domain. (b) Plot of theoretically evaluated growth rate of heterochromatin domains with (red) and without (blue) reactions. Reactions give rise to a stable domain radius. In absence of reactions, there is no stable domain radius.

### S6 Stable domain radius is not significantly regulated by interfacial effects

We have seen via the derivation in Section S3 and Eq S13 that a steady state heterochromatin domain (obtained by setting *dR*/*dt* = 0) is regulated via reaction kinetics, and apparently by the interfacial energy penalty *η*. Here we show that the contribution of *η* in determining 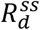 is very small.

**Fig S8:**
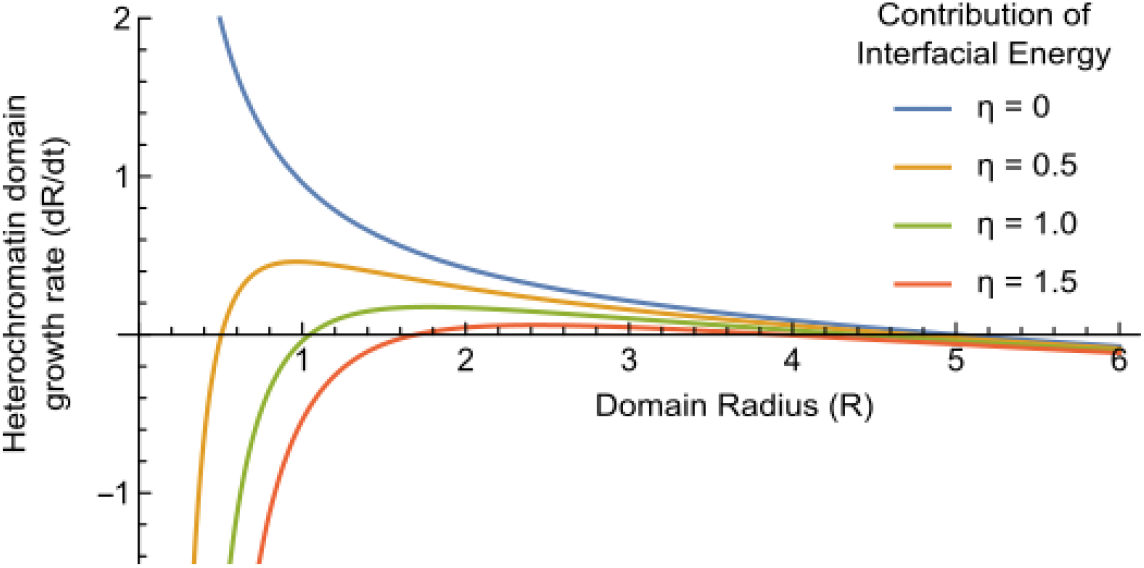
Effect of interfacial energy penalty *η* on the stable radius of heterochromatin domains.

As in the previous section, we plot the growth rate of the heterochromatin droplets with respect to their current radius, as obtained from Eq S13 (Figure S8). The stable radius of the domains 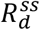 is the point where the curve intersects with the x-axis such that the growth rate of the domains becomes zero, i.e. *dR*/*dt* = 0. As the contribution of the interfacial energy is changed by changing the energetic penalty *η* on the formation of interfaces, we note that the change in the domain sizes is very small. Even when *η* = 0, we see that the domain size does not change appreciably.

These results highlight a key role played by the epigenetic reactions as well as transcriptional regulation of chromatin extrusion. Interestingly, these results also highlight the key difference between our ‘non-equilibrium’ thermodynamic phase-separation model for chromatin organization as opposed to a more traditional energy-minimizing phase-separation. The traditional phase-field models result in domain formation by a competition between the energy reducing phase separation and energy increasing interface formation. However, in our model, the role of the interface formation is overshadowed by the role of reactions in a kinetics rather than energetics driven formation and maintenance of heterochromatin domains. This ‘competition’ between the energetic phase-separation and kinetics of interconverting reactions results in formation of heterochromatin domains of characteristic sizes.

### S7 LAD thickness determination in presence of transcription – theoretical analysis

**Fig S9:**
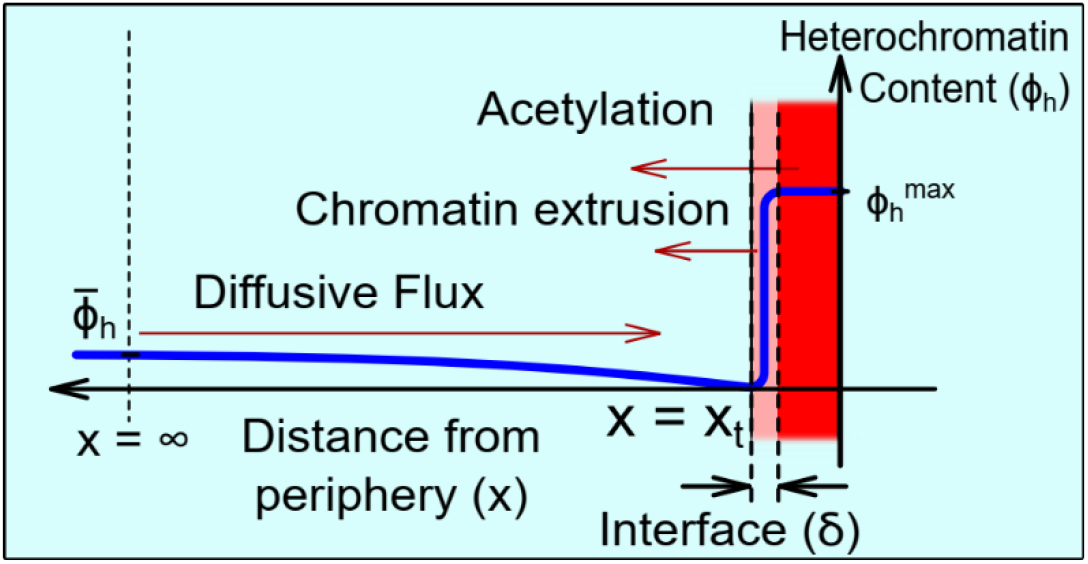
The competition of diffusion driven influx of heterochromatin with the epigenetic reaction and transcription mediated extrusion driven outflux of heterochromatin from the heterochromatin domain determines its steady state size. The figure also shows the radial distribution of heterochromatin volume fraction *ϕ* in and around the domain.

Like the determination of the radius of the heterochromatin droplets in the interior of the nucleus, at the nuclear periphery, the epigenetic, transcriptional and diffusion kinetics balance regulates the thickness of the LADs.

We begin by examining the volume fraction fields within and around the LAD. As discussed in Section S3, the acetylation, methylation and chromatin extrusion together determine the mean heterochromatin (and euchromatin) volume fraction in the nucleus, given by Eq S10. However, a homogenous mean chromatin composition 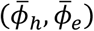 lies in neither of the energy wells as shown in Figure 2b (light blue circle) and is thus energetically unfavorable. Nucleation of heterochromatin domains occurs due to the reduction of free energy as the system evolves. The interaction of heterochromatin with the nuclear lamina results in a preferential nucleation of heterochromatin domains along the lamina i.e lamina associated domains, LADs. We assume that the LADs are formed uniformly along the lamina, and can grow normal to the lamina i.e. increase in thickness, as shown in Figure S9. We also assume that the LADs are far away from the neighboring interior heterochromatin domains, and do not interact with them.

Using a cartesian coordinate system with origin at the nuclear lamina directed normal to it, we determine spatial distribution of the heterochromatin volume fraction *ϕ*_*h*_(*x*). Note that this can be easily done by extending the derivations for the interior heterochromatin domains by setting *R* →

∞, to obtain a linear continuous LAD. Thus,

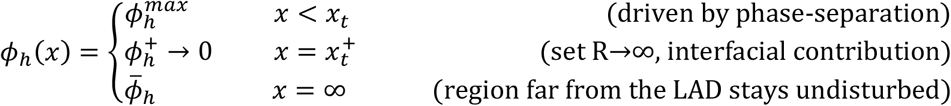

where *x*_*t*_(*t*) is the thickness of the LAD at any time-step. Figure S9 shows the distribution of heterochromatin volume fraction *ϕ*_*h*_(*x*) in the vicinity of a LAD (in red) of thickness *x*_*t*_ as it grows surrounded by euchromatin phase (in blue). The volume fraction field outside the droplet at steady state must follow the evolution equation,

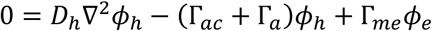

with boundary conditions 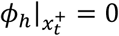 and 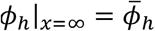, and thus must have the form,

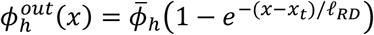

where *ℓ*_*RD*_ is the characteristic reaction-diffusion length scale given under a dilute limit as 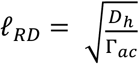. The slope of the heterochromatin profile outside the droplet will drive an inward flux of heterochromatin into the droplet. As for the interior heterochromatin domains, the inward diffusion is opposed by the outward flux of heterochromatin into euchromatin phase which occurs due to both acetylation of histones inside as well as DNA loop extrusion along the domain periphery. Thus, the rate of change of the volume of the droplet *V*_*d*_ can be written as,

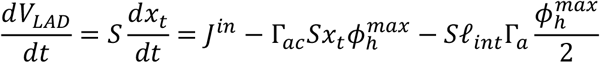

where 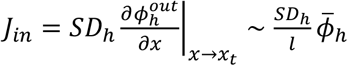 is the diffusive influx. At steady state, by setting *dx*_*t*_/*dt* = 0, we obtain,

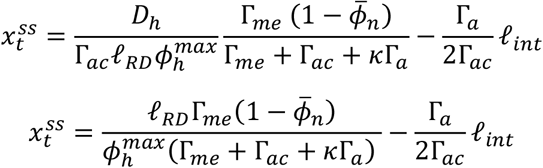

Rescaling,

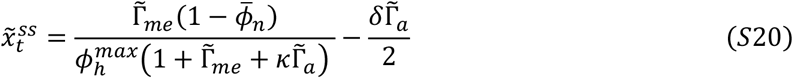

### S8 Model calibration and validation

Having developed the model to capture the spatio-temporal organization of chromatin in the nucleus, we numerically solve Eq S3’ and S8’. The choice of parameters used in our simulations are motivated by the discussions in this section. The parameters can be broadly classified into four types.

#### Kinetic parameters

In the non-dimensional model (Eq S3’ and S8’), all times are rescaled with respect to the timescale of acetylation reaction rate Γ_*ac*_ (as discussed in the extended methods, Section S1), and hence the only parameters which can be altered are the non-dimensional rates of methylation 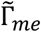 and active chromatin extrusion 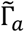. As discussed in section S4, the rates of methylation and acetylation govern the size of the heterochromatin domain and only the acetylation rate determines the intra-domain spacing.

In the rescaled model, the distribution of the heterochromatin domains viz. their sizes is regulated only by 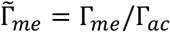. The spacing between the domains is regulated by the size of the domain chosen to model the nucleus. We choose the diameter of nucleus much greater than the reaction-diffusion length-scale (*L*_*nucleus*_ ≫ *ℓ*_*RD*_). Specifically, we chose the nuclear diameter ∼ 20*ℓ*_*RD*_, such that the simulations give the heterochromatin domain spacing qualitatively similar to the inter-domain spacing observed in-vitro via STORM imaging of ActD-treated nuclei. ActD-treated nuclei are specifically chosen for parameter estimation so as to eliminate the effects of transcription mediated chromatin extrusion rate 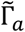 in determining the chromatin organization. Next, the size of the heterochromatin domains is controlled by 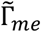. This parameter is obtained quantitatively from the analysis of the images of ActD-treated nuclei so as to get a similar size distribution in the simulation. Also note that the choice of 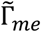 will control the morphology of the heterochromatin domains (discussed in Section S2, Figure S4). We choose 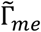 so as to obtain predominantly nearly circular domains so as to facilitate a more straight-forward calculation of heterochromatin domain length-scales. The calibrated values for *L*_*nucleus*_ and 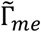 as listed in Table S1. A qualitatively comparable prediction of chromatin organization upon transcription inhibition as well as under control conditions validates our parameter choice (Section 3.3). Note that all the simulations reported in the main manuscript as well as the SI use the same values for *L*_*nucleus*_ and 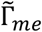. Chromatin extrusion rate Γ_*a*_ is calibrated and validated based on in-vitro nuclear imaging as discussed in Section 3.3 of the main manuscript and algorithmically depicted in Figure S11.

#### Energetic parameters

The non-dimensional energy density (Eq 1 or Eq S1’) involves a single parameter – the chromatin-lamina interaction strength 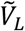. We have previously [14] shown that, 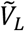 plays a role in deciding the LAD thickness and morphology, i.e. whether the LAD would be more droplet shaped or lamellar. The LAD morphology observed in in-vitro nuclei with transcriptional abrogation is used to estimate the value of 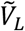. For the previously estimated values of epigenetic reaction rates, we parametrically vary 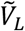 (Figure S10) to obtain a close match with the LAD distribution in ActD treated nucleus (Figure 3d, right panel) as reported in Table S1. All the simulations reported here use the same value for 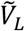, hence validating its choice with the cases where transcription is active.

**Fig S10:**
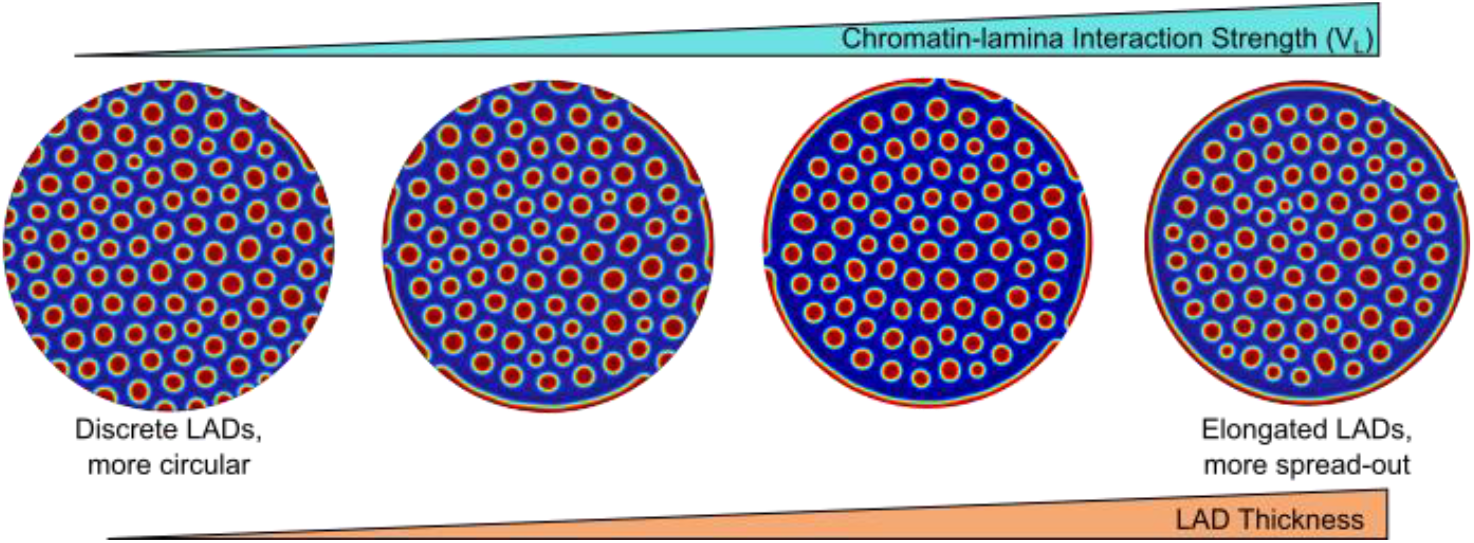
Results of a parametric study on the variation of LAD thickness as the chromatin-lamina interaction strength 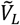 is increased. As 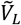 increases, the LADs become more spread out over the entire nuclear periphery. A comparison with distribution of LAD observed in in-vitro nuclei allows the evaluation of the parameter 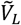. Further note that change in 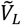 has no effect on the sizes of the heterochromatin domains in the interior of the nucleus.

#### Initial/boundary conditions

We consider an initial spatially homogenous distribution of chromatin and nucleoplasm in the nucleus. The nucleoplasm content of the nucleus is estimated based on experimental images as 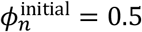, which is maintained a constant in the simulations as there is no exchange exchange of water across the boundary (equivalent to a boundary condition of no flux of nucleoplasm (∇*μ*_*n*_|_boundary_ = 0)). As chromatin is confined to the nucleus, a no flux boundary condition of the order parameter (∇*μ*_*d*_|_boundary_ = 0) ensures the conservation of epigenetic marks.

#### Spatial perturbation parameters

To mimic the spatial heterogeneities of the acetylation and methylation reactions, we add a Gaussian noise with a mean as the parameter values listed in Table SI and a 20% relative standard deviation. This gives us a chromatin domain distribution in agreement with the distribution of domain sizes observed in the experimental images. A similar Gaussian noise is also added to the strength of chromatin-lamina interactions to capture the heterogeneities in the anchoring of chromatin to the lamina. Lastly, we add a random uniform perturbation to the initial chromatin configuration to represent noise due to intrinsic heterogeneities present in the nucleus.

**Table S1:** Values of the simulation parameters

|  | Parameter | Description |  | Value | Remarks |
| --- | --- | --- | --- | --- | --- |
| Initial Conditions | $\phi_n^{initial}$ | Initial water (nucleoplasm) content in the nucleus | | 0.5 | Estimated from experimental H2B density |
| | $\phi_d^{initial}$ | Initial value of order parameter | | 0.1 | The choice does not affect the simulations in presence of reactions |
| | $\sigma_\phi^{noise}$ | Range of uniform perturbation due to heterogeneities in initial conditions | | 0.01 | |
| Epigenetic Kinetics | $\tilde{\Gamma}_{me}$ | Reaction rate of histone methylation (non-dimensionalised) | | 1.2 | Estimated from H2B density in nuclei with transcription inhibited |
| | $\sigma_\Gamma^{noise}$ | Variance in spatial distribution of reaction rates | | 0.2 | Value large enough to obtain a spatial variation in domain sizes |
| Chromatin-lamina Interactions | $\tilde{V}_L$ | Chromatin-lamina interaction strength (non-dimensionalised) | | 0.012 | Parametric study performed to best approximate the LAD distribution in absence of transcription |
| | $\sigma_{V_L}^{noise}$ | Variance in spatial distribution of $\tilde{V}_L$ | | $= \sigma_\Gamma^{noise}$ | |
| Extrusion Kinetics | $\Gamma_a$ | Rate of active chromatin extrusion | Control | 0.03 | Calibrated with experiments |
|  |  |  | ActD treatment | 0 | Total inhibition of loop extrusion |
| | | | WAPLΔ | 0.0075 | Calibrated with experiments |

**Fig S11:**
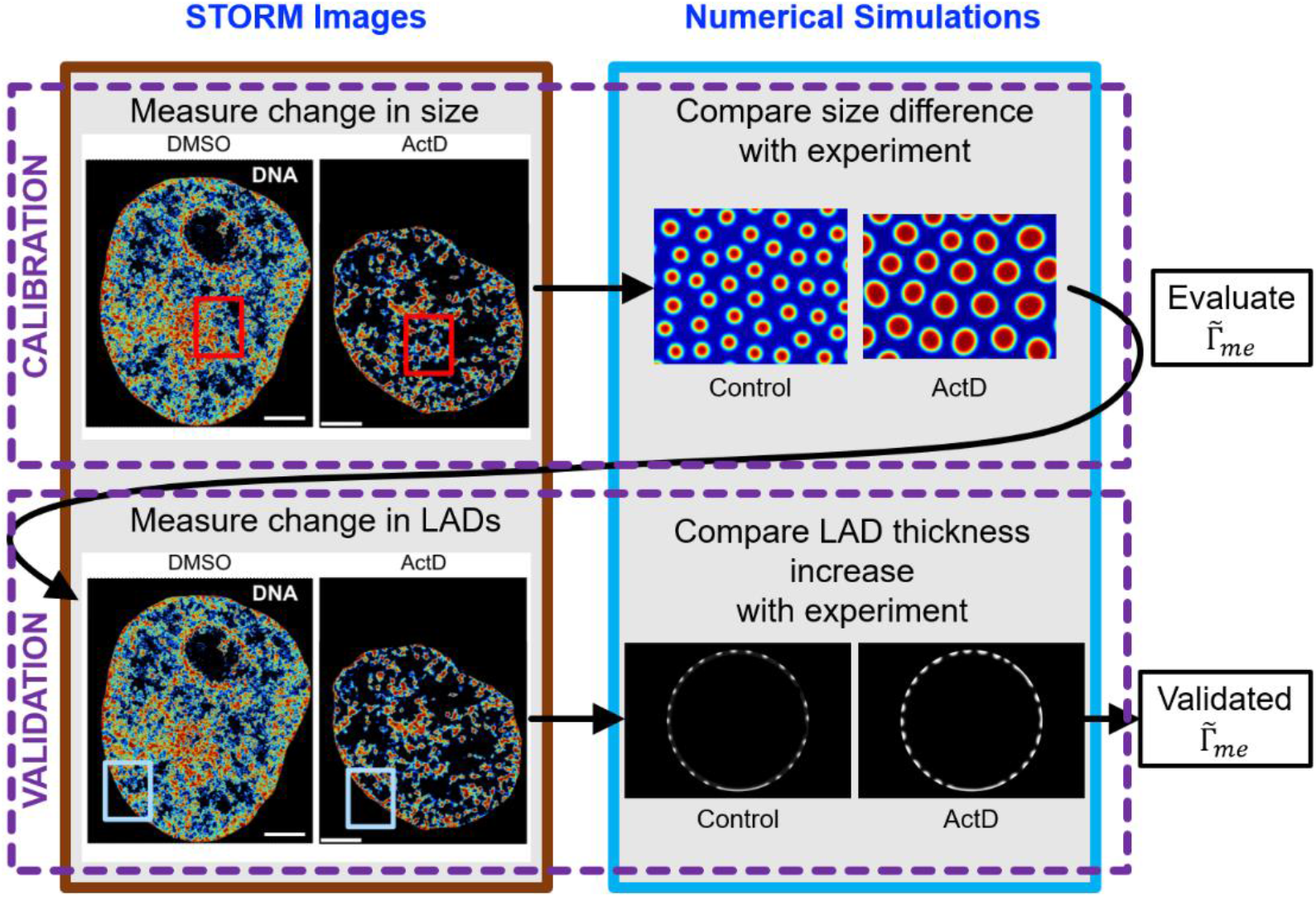
Methodology for calibration and validation of the extrusion rate parameter 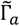 as described in Section 3.3. All scale bars 3 *μ*m.

